# St. Jude Cloud—a Pediatric Cancer Genomic Data Sharing Ecosystem

**DOI:** 10.1101/2020.08.24.264614

**Authors:** Clay McLeod, Alexander M. Gout, Xin Zhou, Delaram Rahbarinia, Andrew Thrasher, Scott Newman, Kirby Birch, Michael Macias, David Finkelstein, Jobin Sunny, Rahul Mudunuri, Brent A. Orr, Madison Treadway, Bob Davidson, Tracy Ard, Andrew Swistak, Stephanie Wiggins, Scott Foy, Samuel W. Brady, Jian Wang, Edgar Sioson, Shuoguo Wang, J. Robert Michael, Yu Liu, Xiaotu Ma, Aman Patel, Michael N. Edmonson, Mark R. Wilkinson, Andrew Frantz, Ti-Cheng Chang, Liqing Tian, Shaohua Lei, Christopher Meyer, Naina Thangaraj, Pamella Tater, Vijay Kandali, Singer Ma, Tuan Nguyen, Omar Serang, Irina McGuire, Nedra Robison, Darrell Gentry, Xing Tang, Lance Palmer, Gang Wu, Ed Suh, Leigh Tanner, James McMurry, Matthew Lear, Zhaoming Wang, Carmen Wilson, Yong Cheng, Mitch Weiss, Gregory T. Armstrong, Leslie L. Robison, Yutaka Yasui, Kim E. Nichols, David W. Ellison, Chitanya Bangur, Charles G. Mullighan, Suzanne J. Baker, Michael Dyer, Geralyn Miller, Michael Rusch, Richard Daly, Keith Perry, James R. Downing, Jinghui Zhang

**Author notes:** Contributed Equally (Clay McLeod, Alexander M. Gout). Correspondence addressed to Keith Perry, James R. Downing and Jinghui Zhang. **STATEMENT OF SIGNIFICANCE** To advance research and treatment of pediatric cancer, we developed St. Jude Cloud, a data sharing ecosystem for accessing >1.2 petabytes of raw genomic data from >10,000 pediatric patients and survivors, innovative analysis workflows, integrative multi-omics visualizations, and a knowledgebase of published data contributed by the global pediatric cancer community.

## Abstract

Effective data sharing is key to accelerating research that will improve the precision of diagnoses, efficacy of treatments and long-term survival of pediatric cancer and other childhood catastrophic diseases. We present St. Jude Cloud (https://www.stjude.cloud), a cloud-based data sharing ecosystem developed via collaboration between St. Jude Children’s Research Hospital, DNAnexus, and Microsoft, for accessing, analyzing and visualizing genomic data from >10,000 pediatric cancer patients, long-term survivors of pediatric cancer and >800 pediatric sickle cell patients. Harmonized genomic data totaling 1.25 petabyes on St. Jude Cloud include 12,104 whole genomes, 7,697 whole exomes and 2,202 transcriptomes, which are freely available to researchers worldwide. The resource is expanding rapidly with regular data uploads from St. Jude’s prospective clinical genomics programs, providing public access as soon as possible rather than holding data back until publication. Three interconnected apps within the St. Jude Cloud ecosystem—Genomics Platform, Pediatric Cancer Knowledgebase (PeCan) and Visualization Community—provide a unique experience for simultaneously performing advanced data analysis in the cloud and enhancing the pediatric cancer knowledgebase. We demonstrate the value of the St. Jude Cloud ecosystem through use cases that classify 48 pediatric cancer subtypes by gene expression profiling and map mutational signatures across 35 subtypes of pediatric cancer.

## INTRODUCTION

Cancer is the number one cause of death by disease among children, with over 15,000 new diagnoses within the United States alone each year (1). The advent of high-throughput genomic profiling technology such as massively parallel sequencing has enabled mapping of the entire 3 billion bases of genetic code for individual human genomes, including those of pediatric cancer. Major pediatric cancer genome research initiatives such as the St. Jude/Washington University Pediatric Cancer Genome Project (PCGP) (2) and NCI’s Therapeutically Applicable Research To Generate Effective Treatments (TARGET, https://ocg.cancer.gov/programs/target) have profiled thousands of pediatric cancer genomes. The resulting data, made accessible through public data repositories such as dbGaP or EGA, have been used to generate new insights into the mechanisms of cancer initiation and progression (3–7), to discover novel targets including those for immunotherapy (8–11), and to build comprehensive genomic landscape maps for developing precision therapy (12–17).

Data sharing, a pre-requisite for genomic research for almost 30 years, is especially important for pediatric cancer, a rare disease with many subtypes driven by diverse and distinct genetic alterations. Based on the annual cancer diagnoses collected from NCI’s Surveillance, Epidemiology and End Results (SEER) program for the period 1990-2016 (18), more than 50% of the pediatric cancer subtypes are rare cancers with an annual incidence of <200 cases in the US (19). Therefore, samples acquired by a single institute, a single research initiative, or, in some instances, even a single nation may lack sufficient power for genomic discovery and clinical correlative analysis. Additionally, the discovery of structural variations and non-coding variants which are important classes of driver variants in pediatric cancer (16,20–22), requires the use of whole-genome sequencing (WGS) to interrogate noncoding regions, which constitute over 98% of the human genome. This imposes another challenge in sharing pediatric cancer genome data as the size of WGS data is ~10 times larger than that of whole-exome sequencing (WES) data which profiles only the coding regions.

To share pediatric cancer genome data using the established public repository model requires major investment in time, professional support and computing resources from users and data providers alike. Under this model (**Fig. 1A**, left), genomic data becomes available for download after submission to a public repository by a computational professional. To use the data, a researcher needs to 1) prepare and submit a request for data access and wait for approval; 2) download data from the public repository to a local computing infrastructure; 3) re-process for data harmonization and annotation using the current reference knowledgebase; 4) perform new analysis or integrative analysis by incorporating custom data; and often 5) submit the new data or the results back to the public repository. With continued expansion of the public data repository and user data, integrating public and local data is an iterative process requiring continued upscaling of local computational resources. Cloud-based technology can establish a shared computing infrastructure for data access and computing for all users, which can improve the efficiency of data analysis by removing the barriers on computational infrastructure required for data transfer and hosting so that computing resources can be dedicated to innovative data analysis and novel methods development (**Fig. 1A**, right).

**Figure 1.**
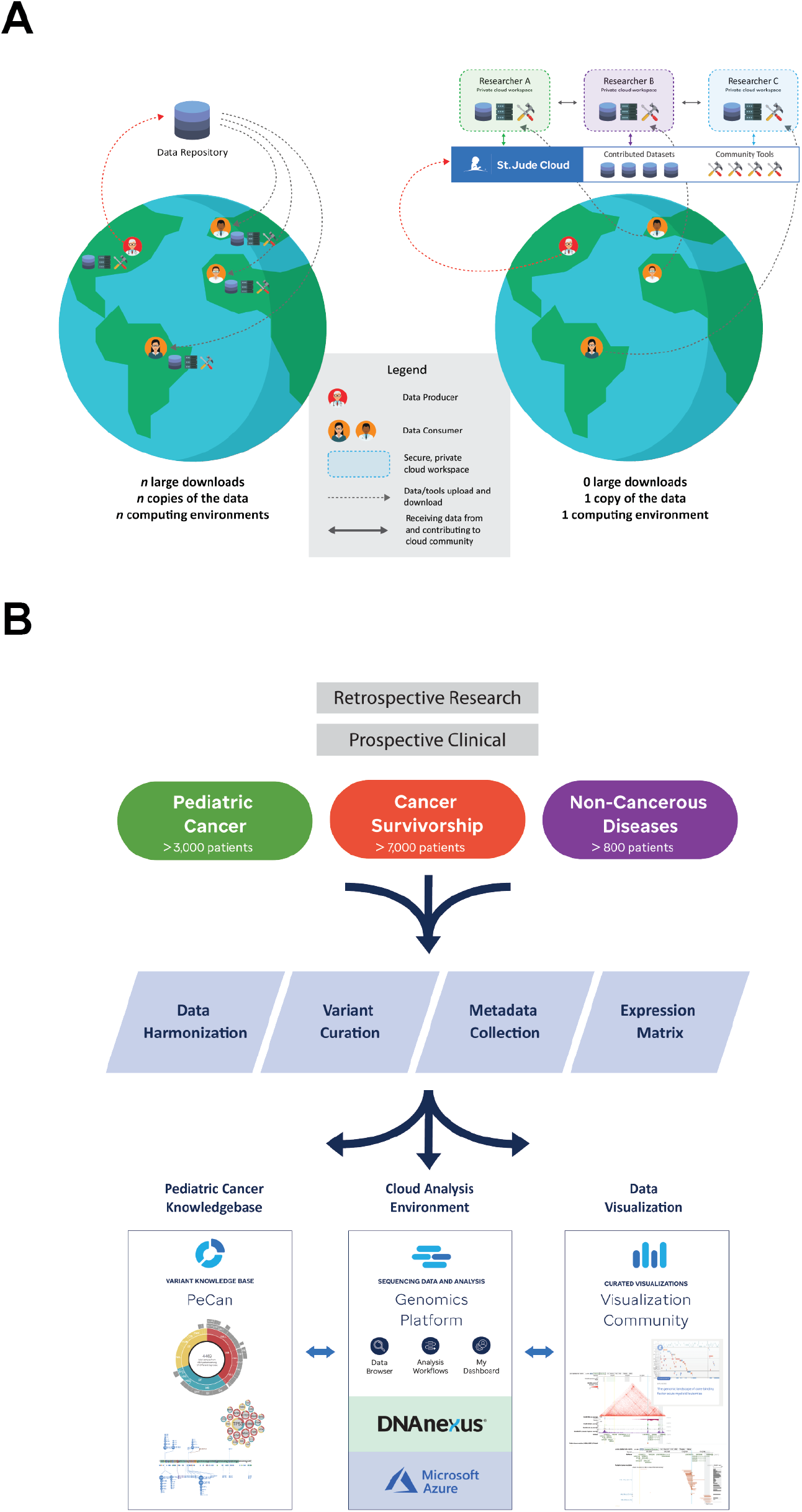
Overview of St. Jude Cloud. (**A**) Comparison of data sharing via the established centralized data repository model versus St. Jude Cloud. The established model requires replication of data and local computing infrastructure while cloud-based data sharing enables a user to perform custom analysis by uploading tools/analysis code onto the shared cloud-computing infrastructure without replication. (**B**) Overview of ingress, harmonization, and deposition of high-throughput sequencing datasets into the St. Jude Cloud ecosystem. Raw genomic data, collected from both retrospective research and prospective clinical studies, were harmonized and curated for access by the broad research community via the three apps on the St. Jude Cloud: Genomics Platform, PeCan Knowledgebase and Visualization Community.

To accelerate research on pediatric cancer and other childhood catastrophic diseases, we developed St. Jude Cloud (https://www.stjude.cloud), a data-sharing ecosystem with open and controlled access to genomic data of >10,000 pediatric cancers generated from both retrospective research projects and prospective clinical genomics programs (**Fig. 1B**) at St. Jude Children’s Research Hospital (St. Jude). St. Jude Cloud was built by St. Jude in partnership with DNAnexus and Microsoft to leverage our combined expertise in pediatric cancer genomic research (2,5,23,24), secure genomic data hosting on the cloud, and Azure cloud computing. St. Jude Cloud is comprised of three interconnected applications: 1) A Genomics Platform that enables controlled access to harmonized raw genomic data as well as end-to-end analysis workflows powered by the innovative algorithms that we developed, tested and validated on data generated from pediatric patient samples; 2) Open access to a knowledgebase portal, PeCan (<u>Pe</u>diatric <u>Can</u>cer), that enables exploration of curated somatic variants of >5,000 pediatric cancer genomes from published literature contributed by St. Jude and other institutions; and 3) A Visualization Community that enables the scientific community to explore published pediatric cancer landscape maps and integrative views of genomic data, epigenetic data and clinical information of pediatric cancers (**Fig. 1B**, bottom). We demonstrate the power of the St. Jude Cloud ecosystem in unveiling important genomic features of pediatric cancer through two use cases: 1) classification of 48 subtypes of pediatric cancer using 1,567 RNA-Seq samples; and 2) characterization of mutational burden and mutation signatures using WGS data generated from 35 subtypes of pediatric cancer.

## RESULTS

### Pediatric Cancer Data Resource on St. Jude Cloud

St. Jude Cloud hosts 12,104 WGS samples, 7,697 WES samples and 2,202 RNA-Seq samples generated from pediatric cancer patients or long-term survivors of pediatric cancer, making it the largest publicly available genomic data resource for pediatric cancer (**Fig. 2A**). Current data sets were acquired from research initiatives such as the St. Jude/Washington University Pediatric Cancer Genome Project (PCGP, (2)), St. Jude Lifetime Cohort Study (SJLIFE, (25)) and Childhood Cancer Survivor Study (CCSS, (26)), as well as from prospective clinical programs such as the Genomes for Kids (G4K) clinical research study of pediatric cancer patients (27) and Real-time Clinical Genomics (RTCG) initiative at St. Jude. Both G4K and RTCG employ a three-platform clinical whole genome, whole exome and transcriptome sequencing of every eligible patient at St. Jude (24). Raw sequence data from all studies were mapped to the latest (hg38) human genome assembly using the same analytical process to ensure data harmonization (Methods). In total, 1.25 petabytes (PB) of genomics data are readily available for access in St. Jude Cloud with over 90% (1.15PB) of this data being WGS data.

**Figure 2.**
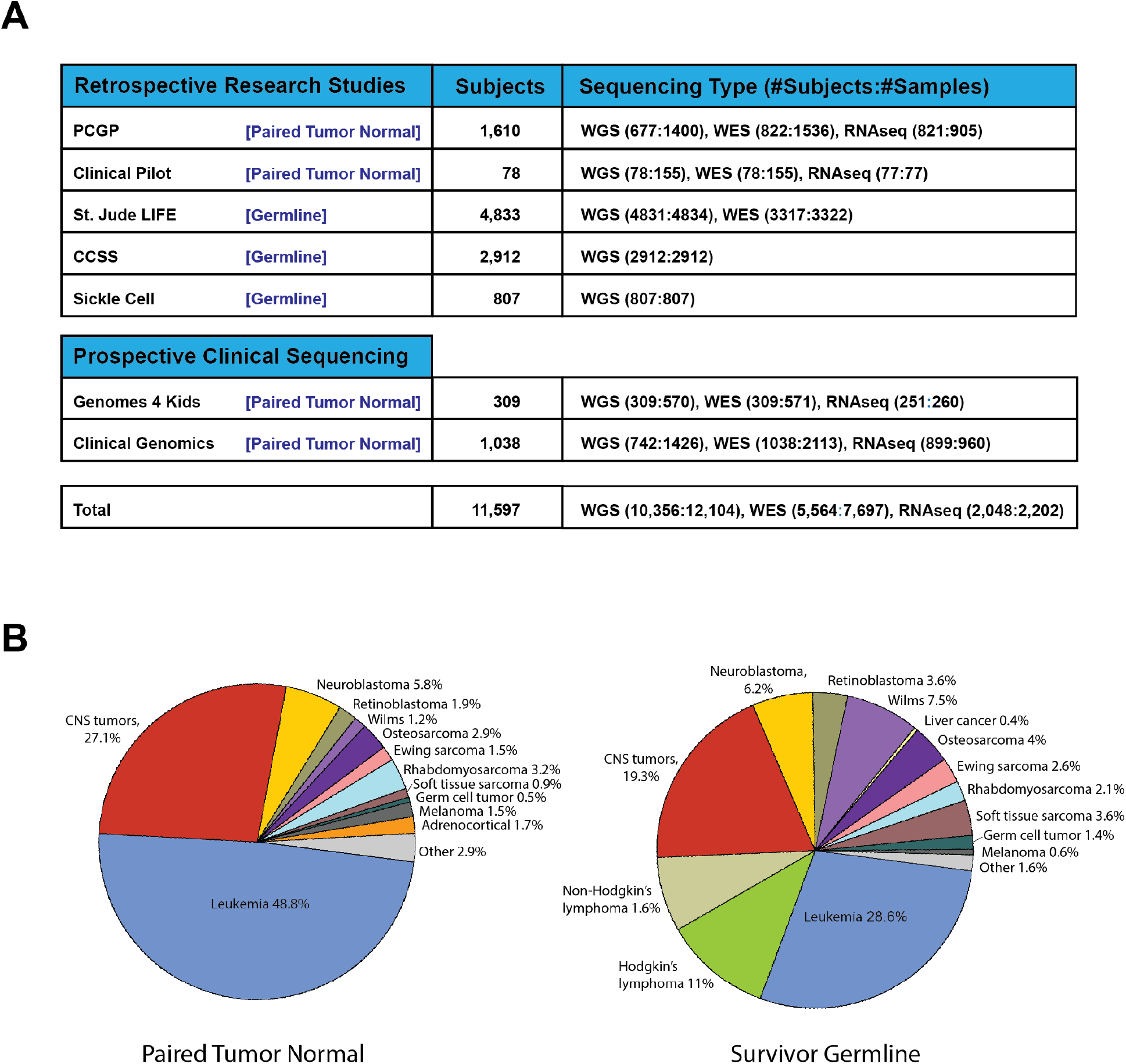
Pediatric cancer genomics data on St. Jude Cloud. (**A**) Summary of high-throughput sequencing data sets on St. Jude Cloud. (**B**) Frequency of pediatric cancer types in WGS data generated from paired tumor-normal samples (left) or germline-only pediatric cancer survivors (right).

When considering only WGS, the collective dataset comprises 3,551 paired tumor-normal pediatric cancer samples and 7,746 germline-only samples of long-term survivors enrolled in SJLIFE or CCSS studies. Major diagnostic categories of the cancer and survivorship genomes, which include pediatric leukemia, lymphoma, CNS tumors and >12 types of non-CNS solid tumors (**Fig. 2B**), are similar except for Hodgkin lymphoma and Non-Hodgkin lymphoma. The lymphoma samples constitute 18% of the cases in the survivorship cohort but are under-represented in the cancer genomes as lymphoma was not selected for pediatric genomic landscape mapping initiatives (*e.g.* PCGP).

### Deposition of Real-time Clinical Genomics (RTCG) Data

As genome-wide sequencing has become an integral part of clinical testing for pediatric cancer, sharing data generated by our CLIA-certified, CAP-accredited clinical laboratory has become an important avenue for expanding the genomic data content on St. Jude Cloud. This allows the data to be made immediately available to investigators at other institutions, rather than held back for months or years until publication. Presently, comprehensive clinical genomic sequencing, including WGS, WES, and RNA-Seq, is offered to all eligible oncology patients at St. Jude as part of their treatment protocols. Many of these patients, or their guardians, consent to sharing their genomic data for research purposes so that it may be utilized to better understand and treat pediatric cancer in the future.

Aiming to share prospective clinical genomic data from appropriately consented patients as quickly as possible, we developed a robust pipeline for the monthly deposition of clinical genomics data generated from the Real-time Clinical Genomics (RTCG) initiative to St. Jude Cloud. As depicted in **Fig. 3A** (details in Methods), the process involves verification of patient consent protocols (and active monitoring for revocation of a previous consent), sample de-identification, remapping to the latest genome build and manual quality checking, all in accordance with legal and ethical guidelines. Basic clinical annotation is retrieved by querying databases of electronic medical records (EMR) and data are harmonized prior to uploading to St. Jude Cloud for public release.

**Figure 3.**
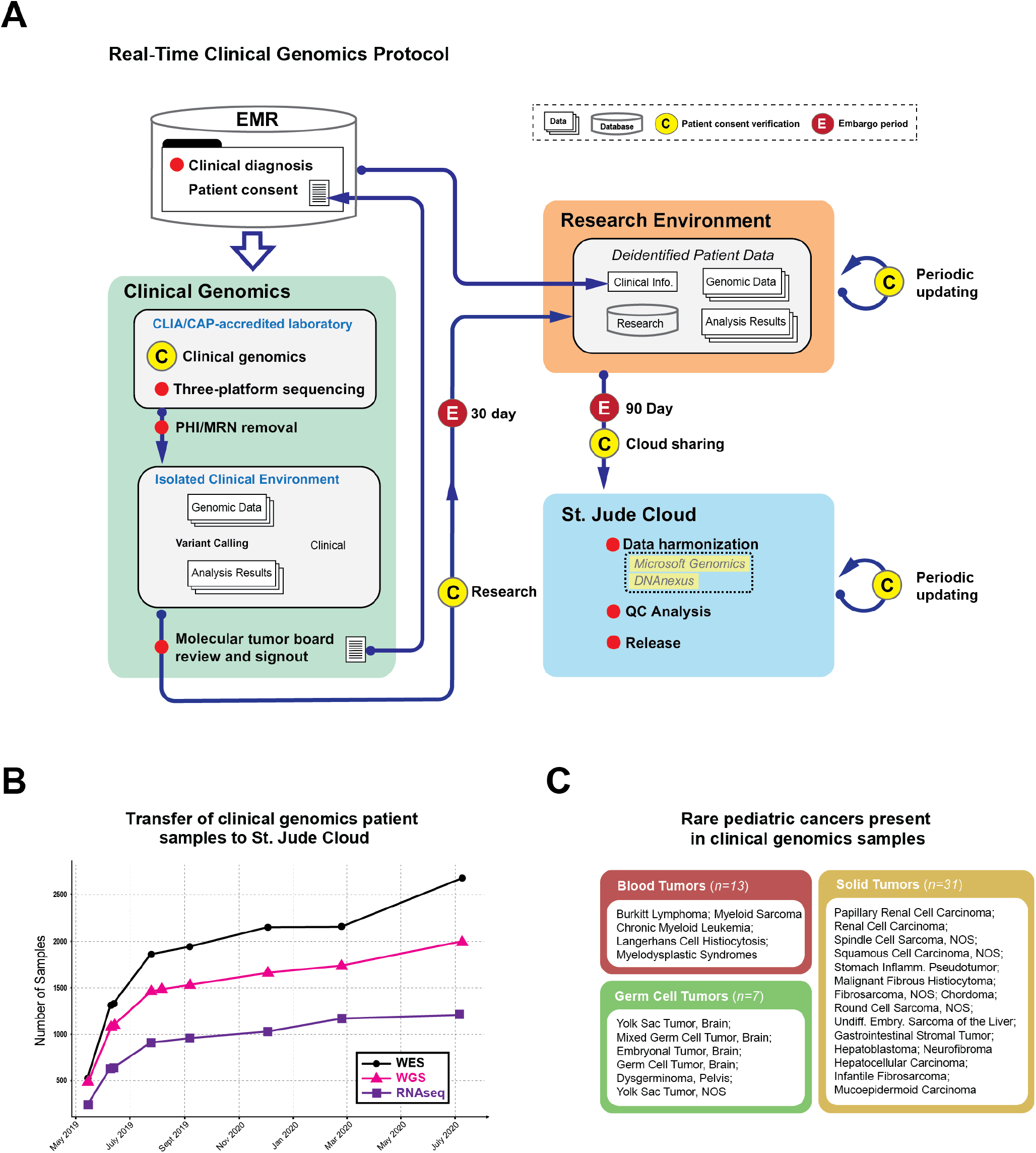
Prospective clinical genomics data on St. Jude Cloud. (**A**) Workflow for monthly deposition of Real-time Clinical Genomics (RTCG) Data. For WGS, WES and RNA-seq data generated from our CLIA/CAP-accredited laboratory, a check for informed consent for research use of patient data is performed following an embargo period (30 days) prior to transfer to the research computing environment. Following a further embargo period (90 days) the genomic data will be uploaded to the St. Jude Cloud with data harmonization performed and data quality assessed prior to release on the Genomics Platform of St. Jude Cloud. (**B**) Cumulative plot of WGS, WES and RNA-seq released on St. Jude Cloud as part of the RTCG deposition process since May 2019. (**C**) Rare pediatric blood (n=13, 5 subtypes), solid (n=31, 16 subtypes) and germ cell (n=7, 6 subtypes) tumor samples uniquely represented in clinical genomics samples.

From March 2019 through July 2020, 1,996 WGS, 2,684 WES and 1,220 RNA-Seq data from prospective pediatric cancer samples were uploaded to St. Jude Cloud (**Fig. 3B**). Importantly, these prospective samples include 51 pediatric cancer samples comprising 27 rare subtypes (**Fig. 3C**) not represented in the retrospective cancer samples on St. Jude Cloud. We anticipate a continued expansion of genomic data on St. Jude Cloud at this pace in the future from prospective samples via RTCG.

### End-to-End Genomic Analysis Workflows

To enable researchers with little to no formal computational training to perform sophisticated genomic analysis, we have deployed end-to-end analysis workflows designed with a point-and-click interface for uploading input files and graphically visualizing the results for scientific interpretation. Advanced computational users can access a command line interface for batch-job submission and run-time parameter optimization. Currently, eight production-grade workflows, tested and used by researchers from St. Jude as well as external institutions, have been deployed on St. Jude Cloud. Comprehensive documentation has been developed and is updated based on user feedback.

Five of these eight workflows have integrated cancer genomic analysis algorithms developed using pediatric cancer data sets such as PCGP; and their performance has been iteratively improved by the growing knowledgebase of pediatric cancer. They include: 1) Rapid RNA-Seq, which predicts gene fusions using the CICERO algorithm (28) that has discovered targetable fusions in high-risk pediatric leukemia (8), high-grade glioma (5) and melanoma (29); 2) PeCanPIE (30), which classifies germline variant pathogenicity using the Medal Ceremony algorithm that was developed to assess germline susceptibility of pediatric cancer (5) and genetic risk for subsequent neoplasms among survivors of childhood cancer (31); 3) NeoeptiopePred, which predicts the immunogenicity of somatic mutations and gene fusions, has characterized the neoepitope landscape of 23 subtypes of pediatric cancer (11); 4) cis-X, which detects non-coding driver variants, and has discovered non-coding drivers in pediatric T-lineage leukemia (32); and 5) SequencErr, which measures and suppresses next-generation sequencing errors (33).

Additionally, we optimized several workflows commonly used by basic research laboratories. These include the 1) ChIP-Seq peak calling pipeline, which detects narrow peaks using MACS2 (34) or broad peaks using SICER (35); 2) WARDEN pipeline, which performs RNA-Seq differential expression using R packages VOOM for normalization and LIMMA for analysis (36); and the 3) Mutational Signature pipeline, which finds COSMIC mutational signatures using a VCF file of somatic SNVs by performing linear modeling using the MutationalPatterns (37) algorithm.

### Pediatric Cancer Knowledgebase (PeCan)

To integrate pediatric cancer genomic data generated by the global research community, we developed PeCan, which assembles somatic variants present at diagnosis or relapse, germline pathogenic variants, and gene expression from the published literature. All data, which is re-annotated and curated to ensure quality and consistency, can be explored dynamically using our visualization tool ProteinPaint (38). Currently, PeCan presents data published by PCGP, TARGET, The German Cancer Research Center, Shanghai Children’s Medical Center, and University of Texas Southwestern Medical Center (**Supp. Table S1**). Variant distribution and expression pattern for a gene of interest can be queried and visualized for 5,161 cancer samples of 23 pediatric cancer subtypes. Curated pathogenic or likely pathogenic variants can also be queried directly and visualized on PeCanPIE’s variant page (30) which presents variant allele frequencies from public databases, results from in-silico prediction and pathogenicity prediction algorithms, related literature, and pathogenicity classification determined by the St. Jude Clinical Genomics tumor board.

### Data Visualization

Data visualization is critical for integrating multi-dimensional cancer genomics data so that researchers can gain insight into the molecular mechanisms that initiate and cause the progression of cancer. We developed generalized tools such as ProteinPaint (38) and GenomePaint (39) that enable dynamic visualization and custom data upload of genomic variants, gene expression, and sample information using either protein or genome as the primary data axis; the user-curated genomic landscape maps for cancer subtypes or pan-cancer studies can also be exported into image files to create figures for multiple scientific publications. Additionally, we developed specialized visualizations to present: a) genome view of chromatin state and gene expression using ChIP-seq and RNA-Seq data generated from mouse/human retina (40) or patient-derived xenografts of pediatric solid tumors (41); b) subgroup clustering using methylation data in medulloblastoma (15) or gene expression data in B-ALL (42); and c) genotype/phenotype correlation for pediatric sickle cell patients and long-term survivors and pediatric cancer (31,43). These expert-curated genomic and epigenomic landscape maps are not only valuable for presenting discoveries in published literature, they can also serve as an important resource for dynamic data exploration by the broad research community.

### St. Jude Cloud Ecosystem

Raw and curated genomic data, analysis and visualization tools are structured into the following three independent and inter-connected applications on St. Jude Cloud to provide a secure, web-based ecosystem for integrative analysis of pediatric cancer genome data: 1) Genomics Platform for accessing data and analysis workflows, 2) PeCan for exploring a curated knowledgebase of pediatric cancer, and 3) Visualization Community for exploring published pediatric cancer genomic or epigenomic landscape maps and for visualizing user data using ProteinPaint or GenomePaint.

A user may work with the St. Jude Cloud ecosystem via open, registered, or controlled access. While PeCan and Visualization Community are accessible in an open and anonymous manner. A user needs to set up a St. Jude Cloud account (register) to run the analysis workflows on the Genomics Platform. In accordance with the community practice for human genomic data protection, access to raw genomic data (*e.g.* WGS, WES or RNA-Seq) generated from patient samples follows a controlled access model, *i.e.* requiring the submission of a signed data access agreement that will be subsequently reviewed by a data access committee for approval. Since its debut in 2018, there are a total of 1,951 registered users of St. Jude Cloud Genomics Platform. 211 requests for access to raw genomic data have been granted to researchers at 80 institutes across 18 countries (**Supp. Figure S1**), and the median turn-around time for data access approval is 7 days. Today, there are ~2,500 unique users per week on average accessing the St. Jude Cloud ecosystem.

While Genomics Platform, PeCan and Visualization Community apps are each valuable resources for pediatric cancer research, working across all three within the St. Jude Cloud ecosystem provides a unique user experience that can simultaneously enhance data analysis and enrich the knowledgebase for pediatric cancer. As illustrated in **Fig. 4**, access to raw genomic data is equivalent to building a virtual research cohort on the St. Jude Cloud ecosystem, which can be accomplished by querying sample features using the data browser of Genomics Platform—a classical approach; or by selecting samples with specific molecular features (*e.g.* mutations or gene expression level) using PeCan. Upon approval, requested data is made available immediately within a private cloud-based project folder. User data can also be uploaded quickly and securely to the project folder through our data transfer tools, and projects can be shared with collaborators using the underlying DNAnexus Platform. The user may then analyze the data using the workflows on the Genomic Platforms, tools provided by the DNAnexus Platform or their own containerized workflows. Alternatively, data can be downloaded to a user’s local computing environment for analysis. Results produced by both local infrastructure or the Genomics Platform can be explored alongside data presented in the curated pediatric cancer knowledgebase (PeCan) using visualization tools such as ProteinPaint or GenomePaint within the Visualization Community. The resulting data, post publication, can be integrated to PeCan to enrich the pediatric cancer knowledgebase, while the landscape maps as well as graphs of sample subgroups prepared by researchers using ProteinPaint or other visualization tools can be shared on the Visualization Community for dynamic exploration. We present two use cases below to demonstrate this process.

**Figure 4.**
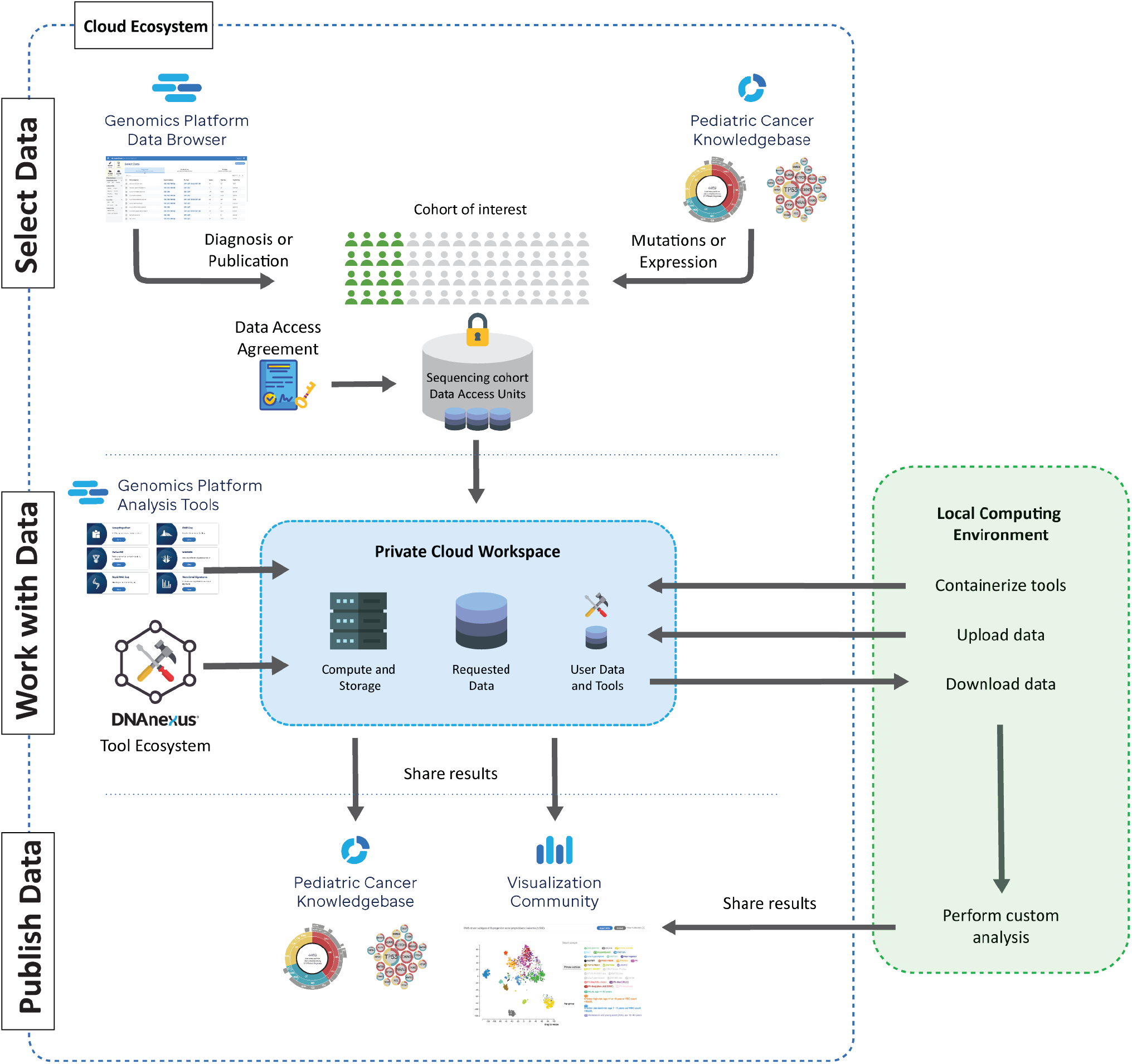
Working across the St. Jude Cloud ecosystem. A virtual cohort can be assembled by querying the data browser on the Genomics Platform (top left) or exploring the Pediatric Cancer Knowledgebase (PeCan) portal (top right). Following approval by the data access committee, the requested data is “vended” onto a private cloud workspace in Genomics Platform (middle center) for analysis using the workflows on St. Jude Cloud (‘Genomics Platform Analysis Tools’), tools available within the DNAnexus Tool Ecosystem, or custom workflows. Alternatively, a user may download the vended genomic data to their local computing infrastructure for further in-depth analysis. Following each of these analyses, a user may share custom visualizations (*e.g.* landscape maps or cancer subgroup analyses) with the research community via the Visualization Community (bottom right) and published results can be incorporated to the PeCan Knowledgebase.

### Use case 1: Expression landscape of pediatric cancers

Defining cancer subtypes by gene expression has provided important insight into the classification of pediatric (44–46) and adult cancers (47). To accomplish this on St. Jude Cloud, we analyzed gene expression profiles of pediatric blood (n=816), solid (n=303) and brain (n=448) tumors using RNA-Seq data from fresh frozen samples which were generated by either retrospective research projects (*e.g.* PCGP and Clinical Pilot) or prospective clinical genomics programs (*e.g.* G4K and RTCG). Gene expression values (**Methods**) were imported from the Genomics Platform and separated into the three categories of blood, solid and brain tumors for subtype classification using t-distributed stochastic neighbor embedding (t-SNE) analysis (**Fig. 5**, https://pecan.stjude.cloud/permalink/stjudecloud-paper.).

**Figure 5.**
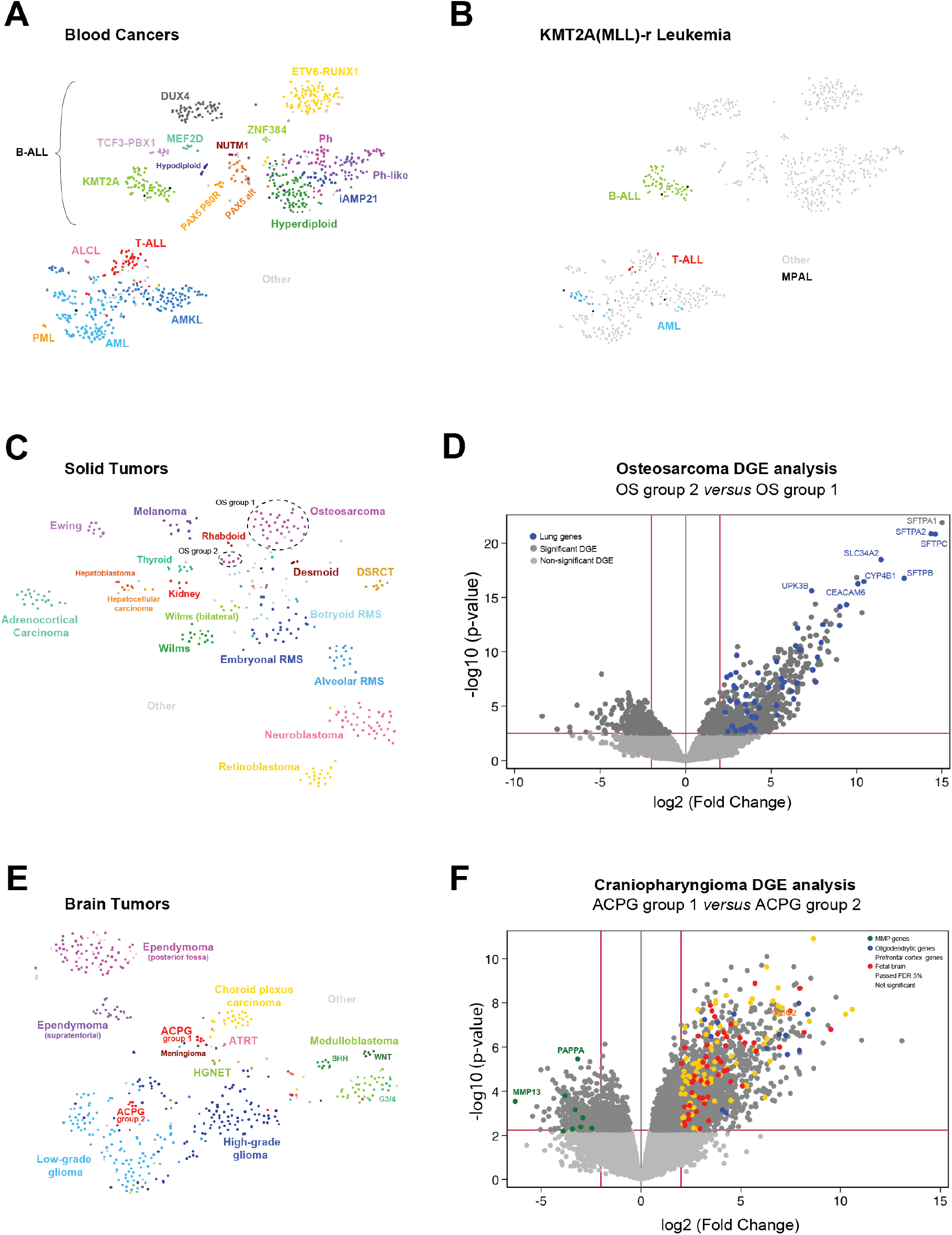
Transcriptome landscape of pediatric cancers. RNA-Seq of 816 blood cancers, 303 solid tumors, and 448 brain tumors were analyzed by a t-distributed stochastic neighbor embedding (t-SNE) algorithm to generate a two-dimensional t-distributed stochastic neighbor embedding (t-SNE) plot highlighting major subtypes in distinct colors. Each dot represents a sample. **(A)** Plot for blood cancers depicting major subtypes comprising B-cell acute lymphoblastic leukemias (B-ALL): *ETV6-RUNX1*, *KMT2A*-rearranged (KMT2A), *TCF3-PBX1*, *DUX4-*rearranged (DUX4), *ZNF384*-rearranged (ZNF384), *MEF2D*-rearranged (MEF2D), *BCR-ABL1* (Ph), Ph-like, Hyperdiploid, Hypodiploid, intrachromosomal amplification of chromosome 21 (iAMP21), *NUTM1*-rearranged (NUTM1), *PAX5* p.Pro80Arg mutation (PAX5 P80R), *PAX5* alterations (PAX5 alt)); T-cell acute lymphoblastic leukemia (T-ALL); acute myeloid leukemia (AML); acute megakaryoblastic leukemia (AMKL), and Anaplastic Large Cell Lymphoma (ALCL). Mixed lineage leukemias harboring KMT2A(MLL)-rearrangements are highlighted in **(B)**. **(C)** Plot for solid tumors displaying major subtypes: osteosarcoma, melanoma, adrenocorticial carcinoma, hepatocellular carcinoma, hepatoblastoma, thyroid papillary tumor (thyroid), kidney cancer, rhabdoid tumor, Ewing sarcoma, Wilms tumor (note bilateral cases form a distinct group), embryononal/alveolar/botryoid rhabdomyosarcoma (RMS), neuroblastoma and retinoblastoma. Two distinct groups of osteosarcoma, denoted OS group 1 and OS group 2 are circled, where the latter are comprised predominantly of metastatic/relapse tumors. **(D)** Volcano plot depicting differential gene expression (DGE) analysis between these two osteosarcoma groups showing an over-representation of Human Gene Atlas ‘Lung’ genes expression (blue) in OS group 2. **(E)** Plot for brain tumors displaying major subtypes: Medulloblastoma (including SHH and WNT subtypes), choroid plexus carcinoma, atypical teratoid/rhabdoid tumor (ATRT), brain germ cell tumor (GCT), meningioma, high-grade neuroepithelial tumor (HGNET), adamantinomatous craniopharyngioma (ACPG), ependymoma (distinguishing posterior fossa versus supratentorial), low-grade glioma, and high-grade glioma. Two observed ACPG groups (1 and 2) are labelled and a **(F)** volcano plot displaying DGE analysis shows an over-representation of oligodendrocytic (blue), prefrontal cortex (yellow), and fetal brain (red) genes in ACPG group 1, while overrepresentation of matrix metallopeptidase (MMP) genes (green) is seen in ACPG group 2.

As shown in **Fig. 5A**, blood cancer expression data reveal a clear distinction between B-cell acute lymphoblastic leukemia (B-ALL, n=521), T-cell acute lymphoblastic leukemia (T-ALL, n=41), acute myeloid leukemia (AML, n=80) and acute megakaryocyte leukemia (AMKL, n=101). The B-ALL subtype clusters here closely recapitulate the subgroups defined by cytogenetic features or gene fusions/somatic mutations reported previously by an analysis of 1,988 childhood or adult B-ALL RNA-Seq (42). The anaplastic large cell lymphoma (ALCL) patient samples, despite their small sample size (n=4 in our cohort), also form a distinct cluster. Interestingly, *KMT2A* (MLL) rearranged leukemias (a subset of which are known to be mixed phenotype acute leukemias) cluster by their cellular lineage (*i.e.* B-cell, T-cell, or myeloid, **Fig. 5B**), indicating their primary lineage has a greater influence than the *KMT2A*-fusion on global gene expression profile.

Solid tumors show tight clusters representing known subtypes (**Fig. 5C**). Interestingly, osteosarcomas cluster into two groups denoted OS group 1 and OS group 2 in **Fig. 5C**, the latter composed of predominantly metastatic tumors. Using the WARDEN pipeline on the Genomic Platform, differential gene expression between the two groups reveals that lung-specific genes are significantly over-represented amongst genes upregulated in OS group 2, with pulmonary-associated surfactant proteins (*SFTPA1*, *SFTAP2*, *SFTPC*, *SFTB*) the most upregulated (**Fig. 5D**, **Supp. Table S2A**). As the OS group2 is comprised of metastatic tumor samples, contamination of the tumor biopsy with lung tissue at the site of metastasis likely contributed to this expression difference. Notably, Wilms tumors also cluster into two distinct groups, one of which is comprised entirely of samples from bilateral cases. This may reflect that divergence in gene transcription caused by different genetic causes of Wilms bilateral versus unilateral cases, likely owing to germline mutations present in the bilateral cases (48).

Brain tumors also form distinct clusters representing major cancer types such as medulloblastoma, choroid plexus carcinoma, ependymoma, craniopharyngioma, high-grade glioma and low-grade glioma (**Fig. 5E**). Consistent with previous reports, medulloblastoma WNT and SHH subtypes are distinct from the larger set of group 3 and 4 subtypes (49) and ependymomas are split into posterior fossa and supratentorial regions (9). Interestingly, adamantinomatous craniopharyngiomas (ACPG), a rare brain cancer derived from pituitary gland embryonic tissue, forms two distinct groups denoted ACPG group 1 and 2 in **Fig. 5E**. WARDEN differential gene expression analysis (**Fig. 5F**) reveals that genes upregulated in ACPG group 1 are significantly over-represented by a wide variety of non-geographically localized neural gene sets including prefrontal cortex, spinal cord, fetal brain and oligodendrocyte (**Supp. Table S2B, C**). By contrast, ACPG group 2 show over-expression of matrix metallopeptidase (MMP) genes (*e.g. MMP13* and *PAPPA*, **Supp. Table S2D**), reported to be associated with cancer progression, migration and metastasis formation (9,50). Examination of genetic and clinical phenotypic data of ACPG group 1 and 2 samples reveals comparable age, gender, frequency of *CTNNB1* mutations (51), and comparable tumor purity ascertained by *CTNNB1* variant allele frequencies (**Supp. Figure 2A**, p-value=0.33), which are correlated with tumor purity (51). Further, both groups displayed high levels of keratin gene expression previously found differentially expressed in ACPG as compared to fetal brain tissue (51) (**Supp. Figure 2B, C**). These observations suggest that the differential gene expression of the two ACPG groups cannot entirely be attributed to differences in tumor purity and possibly indicate a difference in tumor biology. Examination of tumor section slides (**Supp. Figure 2D**) reveals ACPG group 1 tumors display brain invasion and are clearly associated with reactive gliosis while group 2 tumors associate with meninges and cyst walls. This suggests stromal elements are at least partially driving the difference in the transcriptional signature between these two groups of ACPG. Further studies, which may require the use of spatial transcriptomics, are needed to investigate the transcriptional heterogeneity observed in these two groups.

### Use Case 2: Mutation rates and signatures across pediatric blood, solid and brain cancers

Investigation of mutational burden and signatures can unveil the mutational processes shaping the genomic landscape of pediatric cancer (16,17,52) at diagnosis or relapse. To examine mutational burden, we analyzed validated or curated coding and non-coding somatic variants from paired tumor and normal WGS data available for 958 pediatric cancer patient samples comprising over 35 major subtypes of blood, solid, or brain cancers profiled by PCGP, Clinical Pilot or G4K studies (**Fig. 6**, left panel). Among blood cancers, the median number of genome-wide somatic mutation rate was 0.22, 0.31 and 0.37 per million bases (Mb) in AML (including AMKL), B-ALL and T-ALL, respectively. The mutation rate of solid tumors was highly variable by subtype: retinoblastoma had the lowest mutation rate with 0.07 per Mb, while osteosarcoma and melanoma had the highest rates with 1.04 and 8.35 per Mb respectively. Amongst the brain tumors, craniopharyngioma exhibited the lowest mutation rate with 0.02 per Mb in contrast to high-grade gliomas (HGGs) with 0.95 per Mb. Two hypermutators with extremely high mutation burdens were observed among the HGGs owing to mutations in *MSH2* or *POLE*.

**Figure 6.**
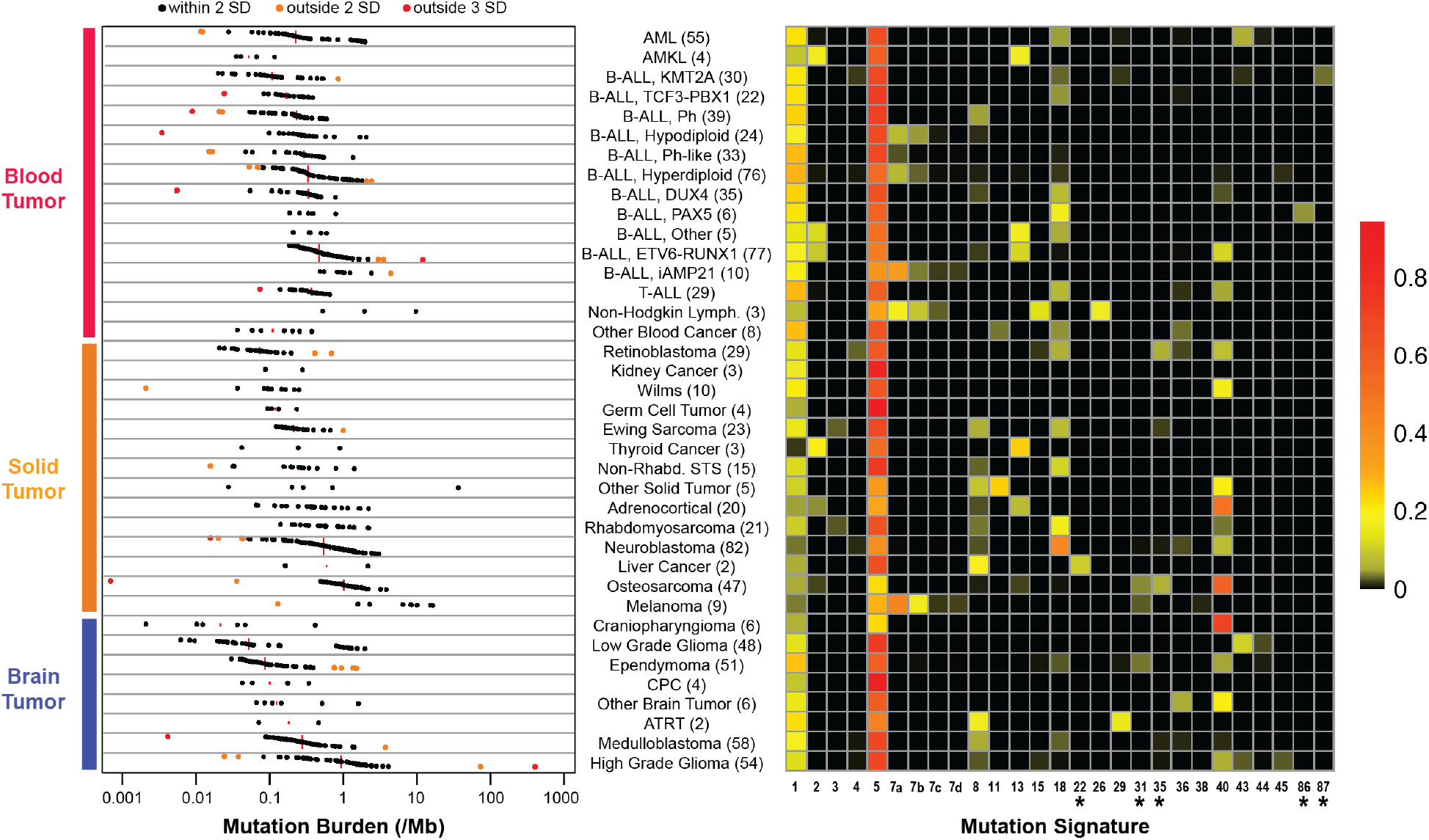
Somatic mutation rate and COSMIC mutation signatures in pediatric cancer subtypes. Both were calculated from coding and non-coding somatic mutations from paired tumor-normal WGS data of blood (samples=456), solid (samples=273) and brain tumor (samples = 229). Mutation burden (left) is shown at a log-scale of number of mutations per Mb. Red lines represent median mutation rate values per subtype. Black, orange, and red indicate samples with mutation rates within two standard deviations (SD), between two and three SD, and greater than three SD within the subtype respectively. Note the outlier osteosarcoma samples with low mutation burden (marked orange and red) have <20% and <10% tumor purity respectively. The orange and red outlier High Grade Glioma samples are hypermutators with bi-allelic loss of either *MSH2* or *POLE*, respectively. Heatmap of COSMIC mutation signatures with therapy-related signatures indicated with an asterisk (*). The scale represents the proportion of somatic mutations contributing to each signature in each sample averaged by subtype. Abbreviations: Acute Myeloid Leukemia (AML), Acute Megakaryocyte Leukemia (AMKL); B-cell acute lymphoblastic leukemia (B-ALL), T-cell Acute Lymphoblastic Leukemia (T-ALL), Non-Rhabdomyosarcoma Soft Tissue Sarcoma (Non-Rhab. STS), Choroid Plexus Carcinoma (CPC), Atypical Teraroid/Rhabdoid Tumor (ATRT). For each subtype the number of samples examined is indicated in parentheses.

First, we performed a preliminary analysis on mutational signatures using the Mutational Signature pipeline on Genomics Platform (**Supp. Figure 3A**) and were able to detect APOBEC mutation signatures (COSMIC signatures 2 and 13) in an acute megakaryoblastic leukemia (AMKL) sample (SJAMLM7006_D in **Supp. Figure 3B**), representing the first implication of this mutagenesis process in AML (52,53). However, spurious signatures may emerge due to potential overfitting. To ensure a more robust analysis, we downloaded somatic VCF files of SNVs detected from the abovementioned paired tumor-normal WGS data set and queried mutational signature abundance using the SigProfilerSingleSample software (23,54), which requires greater stringency to detect signatures. The APOBEC signature in the AMKL sample was reproduced using this approach (**Supp. Figure 3C**).

We detected 25 of the 60 published COSMIC mutation signatures (52) in addition to two recently identified therapy-induced signatures (23) in relapsed B-ALL samples (**Fig. 6**, right). As expected, age-related signatures (*i.e.* COSMIC signature 1 and 5) were present in nearly all pediatric cancers. APOBEC signatures (*i.e.* COSMIC signature 2 and 13) were identified in *ETV6-RUNX1* B-ALL, osteosarcoma, adrenocortical carcinoma, and thyroid cancer, as previously reported (7,53,55,56) and in addition to AMKL as mentioned above. As expected, UV-light induced signature 7 was detected in melanoma and a subset of aneuploid B-ALLs, and, interestingly, in a single case of anaplastic large cell lymphoma (a rare subtype of non-Hodgkin lymphoma). This sample was also positive for signature 15, which is associated with defective DNA mismatch repair. Further, the reactive oxygen species associated signature 18 was found in multiple cancer types including neuroblastoma, rhabdomyosarcoma, T-ALL, Ewing sarcoma, and several subtypes of B-ALL.

Therapy-related signatures were detected in several samples collected post-treatment. The first was signature 22, found in a single hepatoblastoma tumor of an Asian patient that had a mutation rate >10 times higher than the other hepatoblastoma tumors (**Supp. Figure 4**). Interestingly, signature 22 is associated with exposure to aristolochic acid, found in a Chinese medicinal herb (*Aristolochia fangchi*) that is known to be carcinogenic (57). Notably, the relapsed tumor from this patient had increased mutational burden accompanied by acquisition of COSMIC signature 35, which is known to be associated with exposure to cisplatin (**Supp. Figure 4B**), a chemotherapy drug used as part of the standard of care for hepatoblastoma (58). Signature 35 and signature 31, also associated with cisplatin, were found in osteoblastomas and ependymomas as previously reported (59), as well as in retinoblastoma and Ewing’s sarcoma, all of which employ cisplatin for treatment. It is notable that two signatures (currently designated as COSMIC signature 86 and 87) proposed to be induced by ALL treatment were also detected exclusively in relapsed B-ALL samples (**Supp. Figure 5**).

## DISCUSSION

Pediatric cancer is a disease comprised of many rare subtypes. Effective sharing of genomic data and a community effort to elucidate etiology are therefore critical to developing effective therapeutic strategies. St. Jude Cloud is designed to provide a data analysis ecosystem that supports multi-disciplinary research on pediatric cancer by empowering laboratory scientists, clinical researchers, clinicians and bioinformatics scientists. The PeCan portal enables navigation of a pediatric cancer knowledgebase assembled from published literature while the Visualization Community enables dynamic exploration of harmonized and curated data in the forms of landscape maps, cancer subgroups, and integrated views of the genome, transcriptome and epigenome from the same cancer sample. Both apps are designed to be accessible openly by researchers without any formal computational training. Common use cases, such as assessing recurrence of a rare genomic variant or expression status of a gene of interest, are directly enabled by these two St. Jude Cloud apps without the need to download data and perform a custom analysis. If a subset of samples identified through the initial data exploration warrants in-depth investigation, a comprehensive re-analysis can be performed on the Genomics Platform app or a user’s local computing infrastructure. The complementarity amongst the three apps within the St. Jude Cloud ecosystem enables the optimal use of computational resources so that researchers can focus on innovative analysis that will lead to new insight.

User feedback has been critical to informing the trajectory of St. Jude Cloud development. To improve data query, we developed a data browser within the Genomics Platform, which allows a user to select data sets by study, disease subtype, disease stage (*e.g.* diagnosis, relapse or metastasis), sequencing type, and data type. Most recently, RNA-seq feature count data has been made available on the Genomics Platform as these are commonly used for many downstream analyses. We envision an evolving expansion of our current data offerings to include epigenetic and 3D genome data, new facets of our pediatric cancer knowledgebase, non-genomics data, and a variety of additional visualization tools. A new app has been designed for better integration of orthotopic patient-derived xenograft models that are available on the Childhood Solid Tumor Network (CSTN, (41) raw genomic data accessible on the Genomics Platform) and Pediatric Brain Tumor Portal (PBTP, (60), He et al, under review). Moving forward, the rich data resources on St. Jude Cloud may attract external methods developers to use pediatric cancer data—genomic or other data types—as the primary source for development, further expanding the analytical capability of St. Jude Cloud ecosystem and broadening the user base to researchers specializing in other diseases.

A key consideration of our data sharing strategy is to enable access to the pediatric cancer research community as soon as possible, rather than holding data back for publication (which may take months or years). This is accomplished through the development of the Real-Time Clinical Genomics (RTCG) deposition pipeline, a complex workflow involving verification of patient consent, de-identification, data harmonization, and quality checking. To our knowledge, this is the first instance of an institution deposition of prospective clinical genomics data—whole-genome, whole-exome and RNA-Seq—to the scientific research community. The RTCG workflow may serve as a model for other institutions envisioning similar initiatives on sharing data generated from clinical genomics programs with the external community. Currently, the two prospective sequencing projects, RTCG and G4K, have contributed >50% of the raw cancer WGS data on St. Jude Cloud, all of which have been made accessible to 78 investigators from 53 institutions who applied for data access prior to publications of RTCG and G4K. RTCG data has expanded substantially from March to July 2020, at the height of the COVID-19 pandemic in the US (**Fig. 3B**). We anticipate adding approximately 500 additional cases profiled by prospective clinical genomics per year at regular intervals. Data generated from RTCG and G4K are particularly enriched for rare pediatric cancer subtypes (**Fig. 3C**) enabling future research on new therapies that may be incorporated into patient care.

While St. Jude Cloud currently hosts genomic data generated by St. Jude studies, we envision it will serve as a collaborative research platform for the broader pediatric cancer community in the future. User-uploaded data can be analyzed and explored alongside the wealth of curated and raw pediatric genomic data on St. Jude Cloud; because deposition of user data to St. Jude Cloud requires minimal effort. In this regard St. Jude Cloud represents a community resource, framework, and significant contribution to the pediatric genomic sequencing data sharing landscape. We also recognize that contemporary data sharing models are shifting from centralized to distributed resources that serve specific communities. Such distributed repositories are currently not well connected and require considerable effort to move data or tools from one platform to another. The ultimate solution is likely to consist of a federated system for data aggregation, which has also been identified as a priority by participants in the first symposium of The Childhood Cancer Data Initiative (61). This is particularly important for rare subtypes of pediatric cancer as illustrated in our use cases that analyzed craniopharyngioma subgroup classification and hepatoblastoma mutational signature and. An important aspect of future work will be the development of a coordinated effort for data federation across other pediatric genomic resources to enable proper study of these rare tumors.

## METHODS

### St. Jude Cloud Genomics Platform

St. Jude Cloud Genomics Platform is a web application for querying, selecting, and accessing raw and curated genomic data sets through a custom-built Data Browser. Genomic data storage is provided by Microsoft Azure which is accredited to comply with major global security and privacy standards, such as ISO 27001, and has the security and provenance standards required for HIPAA-compliant operation. By leveraging Microsoft Azure, DNAnexus provides an open, flexible and secure cloud platform for St. Jude Cloud to support operational requirements such as the storage and vending of pediatric genomics data to users, along with an environment supportive of genomics analysis tools. DNAnexus supports a security and compliant framework with all of the major data privacy standards (HIPAA, CLIA, CGP, 21 CFR Parts 22, 58, 493, and European data privacy laws and regulations) and interfaces with St. Jude Cloud Genomics Platform. Application for data access can be made using our streamlined electronic process via Docusign (only for requests made within the United States) or a manual process which requires downloading, filling out, signing, and uploading the data access agreement. Upon approval of a data access request by the relevant data access committee(s), St. Jude Cloud Genomics Platform coordinates the provision of a free copy of the requested data to the user via the DNAnexus API into a secure, private workspace within the DNAnexus platform which can also be used for custom data upload.

The Tools section of St. Jude Cloud Genomics Platform provides access to 8 end-to-end St. Jude Cloud workflows optimized for the DNAnexus environment. When a user wishes to run a St. Jude Cloud workflow, St. Jude Cloud Genomics Platform creates a new project folder and vends a copy of the tool to this folder where a user may import St. Jude Cloud genomics data or even upload their own datasets. DNAnexus provides both a command line option for batch execution of operations, and a graphical user-interface for job submission and execution.

### Genomic Sequencing Data

Raw genomic data can be requested and accessed on St. Jude as mapped NGS reads in the BAM (62) file format. The data were generated from paired tumor-normal samples of pediatric cancer patients, germline-only samples of long-term survivors of pediatric cancer, and germline-only samples of pediatric sickle cell patients as summarized in Figure 2A. Paired tumor-normal datasets include retrospective data of 1,610 patients from the St. Jude - Washington University Pediatric Cancer Genome Project (PCGP) (2), 78 patients from a ‘Clinical Pilot’ study (24), and prospective data of 309 patients from ‘Genomes for Kids’ (G4K) study (27) in addition to 1,038 from our real-time clinical genomics (RTCG) initiative. The germline-only dataset of pediatric cancer survivors include 4,833 participants of St. Jude Lifetime Cohort Study (SJLIFE, (25)), a study that brings long-term survivors back to St. Jude Children’s Research Hospital for extensive clinical assessments, and 2,912 participants of the Childhood Cancer Survivor Study (CCSS, (26)), a 31-institution cohort study of long-term survivors. Primary diagnosis of cancer subtypes for both the pediatric cancer and survivorship cohorts is provided both as (i) the value provided at data submission time from the lab or principal investigator (generally unaltered but updated as we receive new information) and as (ii) the harmonized diagnosis value matching the closest classification present in Oncotree (www.oncotree.mskcc.org). Germline-only data of pediatric sickle cell patients include 807 patients from the Sickle Cell Genome Project (SGP), an initiative that is part of the Sickle Cell Clinical Research and Intervention Program (SCCRIP) (63).

Each of these studies represent an individual data access unit within St. Jude Cloud, and was approved for data sharing by the St. Jude Children’s Research Hospital Institutional Review Board (IRB). Further, data is only shared where patient families have consented to research data sharing. For each cohort (*i.e.* pediatric cancer, survivor, or sickle cell), a data access committee has been formed that assess and subsequently approves or rejects data access requests.

### Genomic data harmonization and QC check

WGS and WES data were mapped to hg38 (GRCh38_no_alt) using bwa-mem (64) followed by variant calling using GATK 4.0 HaplotypeCaller (65), both reimplemented by Microsoft Genomics Service (66) on Microsoft Azure, to generate BAM and genomic VCF files for each sample. Each type of genomic sequencing data (WGS, WES) is evaluated separately post-sequencing and mapping. Quality check involves a confirmation of sequence file integrity using Samtools (62) quickcheck and Picard ValidateSamFile (67) and evaluation of the quality, coverage distribution and mapping statistics using Samtools flagstat, FASTQC (68), Qualimap 2 (69) bamqc. The details of the process are described in the respective request for comment (RFC) (70).

RNA-Seq data were mapped to hg38 (GRCh38_no_alt) using a customized workflow (71). In brief, RNA-seq reads were aligned using the STAR aligner in two-pass mode (72) to the human hg38 genome build using gene annotations provided by Gencode v31 gene models (73). Subsequently, Picard (67) SortSam was used to coordinate sort the BAM file, and Picard ValidateSamFile checks that the aligned BAM was valid relative to the format specification. Last, gene-level counts are generated using HTSeq-count (74) using Gencode v31 gene models. For QC check, we used Qualimap 2 RNA-Seq and an in-house “NGSderive strandedness” script (75) that infers strandedness using GENCODE v31 gene annotations.

### Real-Time Clinical Genomics (RTCG) Protocol

Our Institutional Review Board-approved RTCG protocol (St. Jude IRB #19-0099) comprises a series of semi-automated steps that enable the transfer of prospective clinical genomics and selected patient clinical data to St. Jude Cloud. Transfer of this data to St. Jude Cloud is only permitted when patient consent is obtained for clinical genomic testing, research use, and St. Jude Cloud data sharing. This process, depicted in **Fig. 3A**, begins with patient registration and the assignment of PHI/MRN and entry to our electronic medical records database (EMR DB) after which an initial clinical diagnosis is made by the attending physician. Every St. Jude patient has the option of undergoing clinical genomics sequencing as part of our St. Jude clinical genomics service. If patient consent is obtained, the attending physician places an order with the Clinical Genomics team to perform the three-platform sequencing of whole-genome, whole-exome and transcriptome sequencing in CLIA-certified, CAP-accredited laboratory (24). The resulting sequence data is transferred to an isolated clinical computing environment for automated analysis, manual curation, case presentation to our molecular tumor board (MTB) to generate a final case report. Updates to the diagnosis of the patient throughout this process are routine, and we regularly update records based on the most up to date information.

Following the initial MTB sign out of a case report, an embargo period of 30 days is maintained to enable updates or corrections of files prior to the transfer of deidentified genomic data to the research computing environment. Further, clinical information is retrieved from the EMR DB and collated within the research computing environment. After an additional embargo period of 90 days, patient genomic data is transferred to St. Jude Cloud upon verification of consent for cloud data sharing. Once within St. Jude Cloud, data harmonization and QC check are performed as described above prior to public release. Samples are tagged with a rolling publication embargo date which must pass before the data can be used in any external publication. Importantly, patient consent is periodically checked as updates may require the removal of patient data from the research computing and St. Jude Cloud.

### Identification of rare pediatric cancer samples among prospective clinical genomics cohorts

The annual incidence (number of patients per million) of cancer diagnoses (ICCC - International Classification of Childhood Cancer) between the ages of 0-17 years in the USA were calculated using data from the NCI Surveillance, Epidemiology and End Results (SEER) program (19) for the period 1990-2016. Of these, only ICD-O-3 (International Classification of Disease for Oncology, third edition) histology subgroupings with an estimated number of 200 or fewer new patients per million per year were considered rare pediatric cancer subtypes. These estimates were calculated by multiplying the annual incidence per million by 74.2 million, the 2010 census estimate of the number of people in the USA 0-17 years of age. This data was used to determine which of the subtypes unique to the prospective clinical genomics (G4K, RTCG) datasets represented rare cancer subtypes for the St. Jude Cloud platform.

### Pediatric cancer patient sample diagnosis subtype curation

The diagnosis subtype for pediatric cancer patient samples were normalized to a consistent nomenclature across each of the PCGP, Clinical Pilot, G4K, and RTCG sample collections. For PCGP samples, previous associated publications were consulted to ensure accuracy of diagnosis subtype assignment within St. Jude Cloud. For patient samples from Clinical Pilot, G4K and RTCG, clinical genomics pathology reports were used to assign or verify diagnosis subtype annotations. Upon arriving at a concise set of diagnosis subtype annotations across all patient samples on St. Jude Cloud, diagnosis subtype abbreviations were assigned (Supp. Table S3) along with the closest matching Oncotree (www.oncotree.mskcc.org) Identifier.

### Expression analysis of pediatric cancer

St. Jude Cloud tumor RNA-Seq expression count data were generated using HTSeq version 0.11.2 (80) in conjunction with GENCODE (release 31) gene annotations based on the August 2019 release. Of these, only diagnostic, relapse and metastatic samples from fresh frozen tissue (*i.e.* excluding FFPE samples) were included. We removed samples where the associated RNA-seq data involved multiple read lengths or the computationally derived strandedness (InferExperiment (87)) was unclear (samples sequenced using a stranded protocol having less than 80% reverse-oriented stranded read pairs were deemed ‘unclear strandedness’). When patient sample RNA-Seq data was available in both PCGP and Clinical Pilot studies, we only considered the Clinical Pilot data. The analysis only included RNA-seq generated from Illumina GAIIX, HiSeq2000, HiSeq2500, HiSeq4000, NextSeq, or NovaSeq6000 sequencing platforms. These QC steps resulted in a total of qualified 1576 RNA-seq samples which could be queried using the data browser on the St. Jude Cloud Genomics Platform. Once selected, HTSeq feature count files for each of these samples were imported into the St. Jude Cloud ‘*RNA-Seq Expression Classification*’ tool for analysis. In brief, this tool first reads gene features from a GENCODE gene model (release 31). It then aggregates the feature counts from the HTSeq files into a single matrix for all samples under consideration. Covariate information is then retrieved from sample metadata and added to the matrix. Filters are then applied to remove non-protein coding genes and genes exhibiting low expression (<10 read count). This tool also enables subgrouping of samples into ‘blood’, ‘solid’ and ‘brain’ tumor categories (**Supp. Table S3**) of which there were a total of 816, 303, and 448 respective samples (note the sum difference with abovementioned 1576 is from nine germ cell tumors not considered in this analysis). Gene expression analysis was performed with R (3.5.2) using the DESeq2, Rtsne, sva, and plotly packages. Gene expression within each of the blood, solid, and brain, were normalized using DESeq2’s (88) variance stabilizing transformation and batch effects (read length (bp), library strandedness (stranded forward, stranded reverse, and unstranded), RNA selection method (PolyA versus Total RNA), and read pairing (single-versus paired-end)) were removed using ComBat (sva package) (89). The top 1000 most variably expressed genes based on median absolute deviation were then selected from each of the three major cancer types after which two-dimensional t-Distributed Stochastic Neighbor Embedding (t-SNE) was performed according to (42) using a perplexity parameter of 20. Two-dimensional plots for each cancer type were generated using Plotly package.

Differential gene expression analysis for comparison of both osteosarcoma and craniopharyngioma subgroups was performed using the WARDEN pipeline on St. Jude Cloud. Here aligned BAM files were first converted to FASTQ files using bedtools bamtofastq (90). Fastq files were submitted to WARDEN using default parameters. ENRICHR (91,92) was used to perform gene set enrichment analysis using BioGPS Human Gene Atlas, WikiPathways 2019, and GO Molecular Function 2018 gene categories. Volcano plots were generated using STATA/MP 15.1. Adamantinomatous Craniopharyngioma sample tissue section slides were stained with hematoxylin and eosin (H&E stain) and reviewed by a board-certified neuropathologist (BAO).

### Somatic variant data, mutation rate, and mutational signature analysis

Somatic SNVs and indels were analyzed using paired tumor-normal WGS or WES analysis as described previously (24,76). Somatic CNVs were computed using the CONSERTING algorithm (77) followed by manual review of coverage and B-allele fraction. The somatic SNVs/indels and CNVs were lifted over to HG38 and uploaded to St. Jude Cloud as VCF and CNV files.

Mutation rate and signature analysis was performed using all patient tumor sample VCF files from PCGP, Clinical Pilot and G4K studies. When a patient tumor sample VCF file was available in both PCGP and Clinical Pilot studies, we only considered the Clinical Pilot data. The mutation rate was calculated for each subgroup and defined as the number of somatic SNVs per MB. For this purpose, we included only WGS samples and used somatic SNVs in exonic as well as non-exonic, non-repetitive regions (i.e regions not covered by RepeatMasker tracks, sum of these two regions totaling 1,445 Mb).

To identify mutational signatures in these WGS samples, we first determined the trinucleotide context of each somatic SNV using an in-house script, and each sample was summarized based on the number of mutations in each of the 96 possible mutation types (mutation plus trinucleotide context) (53). The presence and strength of 65 COSMIC signatures (52,78) and two therapy-induced mutational signatures which we discovered previously (23) was then analyzed using SigProfilerSingleSample (54) version 1.3 using the default parameters. We selected SigProfilerSingleSample, as it requires greater stringency to prevent overfitting which can lead to spurious signatures. This is accomplished by requiring a cosine increase of 0.05 or above to include a signature, and to include ubiquitous signatures 1 and 5 preferentially prior to detecting additional signatures. Samples explained by the signatures with cosine similarity less than 0.85 were excluded. The proportion of samples (range 0-1) within each cancer subtype category was then displayed in a heatmap to patterns in different cancer subtypes. Mutational signatures within a subtype were only displayed where prevalence exceeds 1%. For the detection of signature 22 in SJST030137, we clustered mutations into diagnosis-specific (present in SJST030137_D1 sample), relapse-specific (present in SJST030137_R1 sample), and shared (present in both samples) clusters, followed by signature analysis with SigProfilerSingleSample on each mutation cluster. The final diagnosis signature spectrum was achieved by summing the signatures in the diagnosis-specific and shared mutation clusters, while the relapse spectrum was the sum of the relapse-specific and shared clusters. This increased sensitivity of detection of signature 22, which was otherwise obscured in the relapse sample due to an increased mutation burden associated with the cisplatin signature.

## DATA AVAILABILITY

All data is available on St. Jude Cloud (www.stjude.cloud). Interactive t-SNE RNA-Seq expression maps are available as a collection within the St. Jude Cloud Visualization Community here: https://viz.stjude.cloud/stjudecloud/collection/stjudecloud-paper. RNA-Seq derived HTSeq count data for samples considered in *Use Case 1: Expression landscape of pediatric cancers*, and somatic VCF files used for mutation burden and mutation signatures analysis in *Use Case 2:Mutation rates and signatures across pediatric blood, solid and brain cancers*, can be accessed through the St. Jude Cloud platform data browser here: https://pecan.stjude.cloud/permalink/stjudecloud-paper

## AUTHOR CONTRIBUTIONS

J.Z., J.R.D., and K.P. conceived the project; J.Z., Clay McLeod, M.R., S.N., and K.B. designed St. Jude Cloud ecosystem; X.Z. developed the visualization tools and A.M.G. led the data analysis along with A.T., D.F., S.N., S.W.B., under the supervision of J.Z., and B.A.O. analyzed clinical data for adamantinomatous craniopharyngioma. K.B., M.T. and D.F. provided user support. Clay McLeod, D.R., M.M., J.S., R.M., B.D., T. A., A.S., S.W., S.F., J.W., E.S., S.W., J.R.M., M.R.W., A.F., S. L., Christopher Meyer, N.T., P.T., V.K., S.M., T.N., O.S., I.M., N.R., D.G., G.W., E.S., L.T., J.M., S.L., A.M.G., and C.B. developed software and/or performed data harmonization under the supervision of J.Z., K.P., Clay McLeod, M.R., C.B., G.M and R.D. Clay McLeod, S.N., M.M., J.S., A.F., Y.L., X.T., L.P., Y.C., T.-C.C., X.M., A.P., M.N.E., L.T., A.T., and A.M.G. developed the analysis workflows. A.M.G. led the cancer subtype diagnosis harmonization along with S.N., Clay McLeod, S.F., D.R., D.F. and R.M. J.R.D., C.G.M, S.J.B. and M.D. contributed the PCGP data; J.R.D., K.E.M., C.G.M, M.L., and D.W.E. contributed the G4K and RTCG data; Z.W., C.W., L.L.R., Y.Y. contributed St. Jude Life data; G.T.A. contributed the CCSS data; and M.W. contributed the sickle cell genomic data. J.Z., A.M.G., and Clay McLeod wrote the manuscript with critical feedback from M.N.E., Y.L., L.L.R., and S.W.B.

## ACKNOWLEDGEMENTS

We wish to thank all St. Jude patients and their families. We would like to thank the generous support from Microsoft AI for Good program for providing Microsoft Azure as the cloud storage for St. Jude Cloud and for supplying Microsoft Genomics Service for WGS and WES harmonization. We would like to thank Mr. Kevin Rodell and Mr. Judson Althoff of Microsoft for initiating the St. Jude/Microsoft Collaboration. We would also like to thank the generous support of DNAnexus. We would like to acknowledge the contribution by members of the St. Jude Biorepository and Clinical Genomics teams for their assistance in developing the RTCG pipeline. We would like to thank: Katherine Steuer for her assistance in verifying patient consent on research data sharing; Dr. Alberto Pappo for consultation on treatment protocols for pediatric hepatoblastoma patients; Drs. David Wheeler, Jennifer Neary, and Antonina Silkov for their help in curating the sample information of RTCG and the analysis of adamantinomatous craniopharyngioma (ACPG) samples; Dr. Diane Flasch for critical review of the manuscript and Drs. Tanja Gruber and Anna Hagstrom for assistance with the clarification of the lineages of MLL-rearranged infant ALL. We like to thank all the users who have provided critical feedback, in particular Drs. Jackie Norrie, Lawryn Kasper, and Laura Hover. This work is funded as a St. Jude Blue Sky initiative.

## Notes

**CONFLICTS OF INTEREST** C.G.M. has received research funding from Abbvie and Pfizer and served on an advisory board for Illumina. Christopher Meyer, N.T., S.A., T.N., O.S., and T.A. are employees of DNAnexus. B.D., T.A., R.I.D., C.B., G.M., and R.D. are employees of Microsoft.

https://www.stjude.cloud/

### Competing Interest Statement

C.G.M. has received research funding from Abbvie and Pfizer and served on an advisory board for Illumina.
Christopher Meyer, N.T., S.A., T.N., O.S., and T.A. are employees of DNAnexus. 
B.D., T.A., R.I.D., C.B., G.M., and R.D. are employees of Microsoft.

## REFERENCES

1. Cunningham RM, Walton MA, Carter PM. The Major Causes of Death in Children and Adolescents in the United States. N Engl J Med 2018;379(25):2468–75 doi 10.1056/NEJMsr1804754.

2. Downing JR, Wilson RK, Zhang J, Mardis ER, Pui CH, Ding L, et al. The Pediatric Cancer Genome Project. Nat Genet 2012;44(6):619–22 doi 10.1038/ng.2287.

3. Zhang J, Benavente CA, McEvoy J, Flores-Otero J, Ding L, Chen X, et al. A novel retinoblastoma therapy from genomic and epigenetic analyses. Nature 2012;481(7381):329–34 doi 10.1038/nature10733.

4. Wu G, Broniscer A, McEachron TA, Lu C, Paugh BS, Becksfort J, et al. Somatic histone H3 alterations in pediatric diffuse intrinsic pontine gliomas and non-brainstem glioblastomas. Nat Genet 2012;44(3):251–3 doi 10.1038/ng.1102.

5. Zhang J, Walsh MF, Wu G, Edmonson MN, Gruber TA, Easton J, et al. Germline Mutations in Predisposition Genes in Pediatric Cancer. N Engl J Med 2015;373(24):2336–46 doi 10.1056/NEJMoa1508054.

6. Ma X, Edmonson M, Yergeau D, Muzny DM, Hampton OA, Rusch M, et al. Rise and fall of subclones from diagnosis to relapse in pediatric B-acute lymphoblastic leukaemia. Nat Commun 2015;6:6604 doi 10.1038/ncomms7604.

7. Brady SW, Ma X, Bahrami A, Satas G, Wu G, Newman S, et al. The Clonal Evolution of Metastatic Osteosarcoma as Shaped by Cisplatin Treatment. Mol Cancer Res 2019;17(4):895–906 doi 10.1158/1541-7786.MCR-18-0620.

8. Roberts KG, Li Y, Payne-Turner D, Harvey RC, Yang YL, Pei D, et al. Targetable kinase-activating lesions in Ph-like acute lymphoblastic leukemia. N Engl J Med 2014;371(11):1005–15 doi 10.1056/NEJMoa1403088.

9. Parker M, Mohankumar KM, Punchihewa C, Weinlich R, Dalton JD, Li Y, et al. C11orf95-RELA fusions drive oncogenic NF-kappaB signalling in ependymoma. Nature 2014;506(7489):451–5 doi 10.1038/nature13109.

10. Zhang J, Wu G, Miller CP, Tatevossian RG, Dalton JD, Tang B, et al. Whole-genome sequencing identifies genetic alterations in pediatric low-grade gliomas. Nat Genet 2013;45(6):602–12 doi 10.1038/ng.2611.

11. Chang TC, Carter RA, Li Y, Li Y, Wang H, Edmonson MN, et al. The neoepitope landscape in pediatric cancers. Genome Med 2017;9(1):78 doi 10.1186/s13073-017-0468-3.

12. Wu G, Diaz AK, Paugh BS, Rankin SL, Ju B, Li Y, et al. The genomic landscape of diffuse intrinsic pontine glioma and pediatric non-brainstem high-grade glioma. Nat Genet 2014;46(5):444–50 doi 10.1038/ng.2938.

13. Tirode F, Surdez D, Ma X, Parker M, Le Deley MC, Bahrami A, et al. Genomic landscape of Ewing sarcoma defines an aggressive subtype with co-association of STAG2 and TP53 mutations. Cancer Discov 2014;4(11):1342–53 doi 10.1158/2159-8290.CD-14-0622.

14. Liu Y, Easton J, Shao Y, Maciaszek J, Wang Z, Wilkinson MR, et al. The genomic landscape of pediatric and young adult T-lineage acute lymphoblastic leukemia. Nat Genet 2017;49(8):1211–8 doi 10.1038/ng.3909.

15. Northcott PA, Buchhalter I, Morrissy AS, Hovestadt V, Weischenfeldt J, Ehrenberger T, et al. The whole-genome landscape of medulloblastoma subtypes. Nature 2017;547(7663):311–7 doi 10.1038/nature22973.

16. Ma X, Liu Y, Liu Y, Alexandrov LB, Edmonson MN, Gawad C, et al. Pan-cancer genome and transcriptome analyses of 1,699 paediatric leukaemias and solid tumours. Nature 2018;555(7696):371–6 doi 10.1038/nature25795.

17. Grobner SN, Worst BC, Weischenfeldt J, Buchhalter I, Kleinheinz K, Rudneva VA, et al. The landscape of genomic alterations across childhood cancers. Nature 2018;555(7696):321–7 doi 10.1038/nature25480.

18. Surveillance, Epidemiology, and End Results Program. NCI <https://seer.cancer.gov/statistics/>. Accessed 2018.

19. Howlader N, A. M. Noone, M. Krapcho, D. Miller, A. Brest, M. Yu, J. Ruhl, Z. Tatalovich, A. Mariotto, D. R. Lewis, H. S. Chen, E. J. Feuer, and K. A. Cronin. 2019 SEER Cancer Statistics Review, 1975-2016. In Based on November 2018 SEER data submission, posted to the SEER web site. National Cancer Institute <https://seer.cancer.gov/csr/1975_2016/ >. Accessed 2020.

20. Mansour MR, Abraham BJ, Anders L, Berezovskaya A, Gutierrez A, Durbin AD, et al. Oncogene regulation. An oncogenic super-enhancer formed through somatic mutation of a noncoding intergenic element. Science 2014;346(6215):1373–7 doi 10.1126/science.1259037.

21. Northcott PA, Lee C, Zichner T, Stutz AM, Erkek S, Kawauchi D, et al. Enhancer hijacking activates GFI1 family oncogenes in medulloblastoma. Nature 2014;511(7510):428–34 doi 10.1038/nature13379.

22. Zimmerman MW, Liu Y, He S, Durbin AD, Abraham BJ, Easton J, et al. MYC Drives a Subset of High-Risk Pediatric Neuroblastomas and Is Activated through Mechanisms Including Enhancer Hijacking and Focal Enhancer Amplification. Cancer Discov 2018;8(3):320–35 doi 10.1158/2159-8290.CD-17-0993.

23. Li B, Brady SW, Ma X, Shen S, Zhang Y, Li Y, et al. Therapy-induced mutations drive the genomic landscape of relapsed acute lymphoblastic leukemia. Blood 2020;135(1):41–55 doi 10.1182/blood.2019002220.

24. Rusch M, Nakitandwe J, Shurtleff S, Newman S, Zhang Z, Edmonson MN, et al. Clinical cancer genomic profiling by three-platform sequencing of whole genome, whole exome and transcriptome. Nat Commun 2018;9(1):3962 doi 10.1038/s41467-018-06485-7.

25. Hudson MM, Ehrhardt MJ, Bhakta N, Baassiri M, Eissa H, Chemaitilly W, et al. Approach for Classification and Severity Grading of Long-term and Late-Onset Health Events among Childhood Cancer Survivors in the St. Jude Lifetime Cohort. Cancer Epidemiol Biomarkers Prev 2017;26(5):666–74 doi 10.1158/1055-9965.EPI-16-0812.

26. Robison LL, Armstrong GT, Boice JD, Chow EJ, Davies SM, Donaldson SS, et al. The Childhood Cancer Survivor Study: a National Cancer Institute-supported resource for outcome and intervention research. J Clin Oncol 2009;27(14):2308–18 doi 10.1200/JCO.2009.22.3339.

27. Genome4Kids. <https://clinicaltrials.gov/ct2/show/NCT02530658>.

28. Tian L, Li Y, Edmonson MN, Zhou X, Newman S, McLeod C, et al. CICERO: a versatile method for detecting complex and diverse driver fusions using cancer RNA sequencing data. Genome Biol 2020;21(1):126 doi 10.1186/s13059-020-02043-x.

29. Newman S, Fan L, Pribnow A, Silkov A, Rice SV, Lee S, et al. Clinical genome sequencing uncovers potentially targetable truncations and fusions of MAP3K8 in spitzoid and other melanomas. Nat Med 2019;25(4):597–602 doi 10.1038/s41591-019-0373-y.

30. Edmonson MN, Patel AN, Hedges DJ, Wang Z, Rampersaud E, Kesserwan CA, et al. Pediatric Cancer Variant Pathogenicity Information Exchange (PeCanPIE): a cloud-based platform for curating and classifying germline variants. Genome Res 2019;29(9):1555–65 doi 10.1101/gr.250357.119.

31. Wang Z, Wilson CL, Easton J, Thrasher A, Mulder H, Liu Q, et al. Genetic Risk for Subsequent Neoplasms Among Long-Term Survivors of Childhood Cancer. J Clin Oncol 2018;36(20):2078–87 doi 10.1200/JCO.2018.77.8589.

32. Liu Y, Li C, Shen S, Chen X, Szlachta K, Edmonson MN, et al. Discovery of regulatory noncoding variants in individual cancer genomes by using cis-X. Nat Genet 2020 doi 10.1038/s41588-020-0659-5.

33. Ma X, Shao Y, Tian L, Flasch DA, Mulder HL, Edmonson MN, et al. Analysis of error profiles in deep next-generation sequencing data. Genome Biol 2019;20(1):50 doi 10.1186/s13059-019-1659-6.

34. Zhang Y, Liu T, Meyer CA, Eeckhoute J, Johnson DS, Bernstein BE, et al. Model-based analysis of ChIP-Seq (MACS). Genome Biol 2008;9(9):R137 doi 10.1186/gb-2008-9-9-r137.

35. Xu S, Grullon S, Ge K, Peng W. Spatial clustering for identification of ChIP-enriched regions (SICER) to map regions of histone methylation patterns in embryonic stem cells. Methods Mol Biol 2014;1150:97–111 doi 10.1007/978-1-4939-0512-6_5.

36. Law CW, Chen Y, Shi W, Smyth GK. voom: Precision weights unlock linear model analysis tools for RNA-seq read counts. Genome Biol 2014;15(2):R29 doi 10.1186/gb-2014-15-2-r29.

37. Blokzijl F, Janssen R, van Boxtel R, Cuppen E. MutationalPatterns: comprehensive genome-wide analysis of mutational processes. Genome Med 2018;10(1):33 doi 10.1186/s13073-018-0539-0.

38. Zhou X, Edmonson MN, Wilkinson MR, Patel A, Wu G, Liu Y, et al. Exploring genomic alteration in pediatric cancer using ProteinPaint. Nat Genet 2016;48(1):4–6 doi 10.1038/ng.3466.

39. Zhuo X. 2020 <https://genomepaint.stjude.cloud/>.

40. Wang L, Hiler D, Xu B, AlDiri I, Chen X, Zhou X, et al. Retinal Cell Type DNA Methylation and Histone Modifications Predict Reprogramming Efficiency and Retinogenesis in 3D Organoid Cultures. Cell Rep 2018;22(10):2601–14 doi 10.1016/j.celrep.2018.01.075.

41. Stewart E, Federico SM, Chen X, Shelat AA, Bradley C, Gordon B, et al. Orthotopic patient-derived xenografts of paediatric solid tumours. Nature 2017;549(7670):96–100 doi 10.1038/nature23647.

42. Gu Z, Churchman ML, Roberts KG, Moore I, Zhou X, Nakitandwe J, et al. PAX5-driven subtypes of B-progenitor acute lymphoblastic leukemia. Nat Genet 2019;51(2):296–307 doi 10.1038/s41588-018-0315-5.

43. Lance E. Palmer XZ, Clay McLeod, Evadnie Rampersaud, Jeremie H. Estepp, Xing Tang, Jian Wang, Edgar Siosan, J. Robert Michael, Kirby Birch, Jason R Hodges, Martha Villavicencio, Michael Rusch, Scott Newman, Heather Mulder, John Easton, Keith Perry, James R. Downing, MD, Jane S. Hankins, Gang Wu, Jinghui Zhang, Mitchell J. Weiss. Data Access and Interactive Visualization of Whole Genome Sequence of Sickle Cell Patients within the St. Jude Cloud. . ASH Annual Meeting. Volume 32 San Diego: Blood; 2018. p 723.

44. Golub TR, Slonim DK, Tamayo P, Huard C, Gaasenbeek M, Mesirov JP, et al. Molecular classification of cancer: class discovery and class prediction by gene expression monitoring. Science 1999;286(5439):531–7 doi 10.1126/science.286.5439.531.

45. Downing JR. Acute leukemia: subtype discovery and prediction of outcome by gene expression profiling. Verh Dtsch Ges Pathol 2003;87:66–71.

46. Kohlmann A, Bullinger L, Thiede C, Schaich M, Schnittger S, Dohner K, et al. Gene expression profiling in AML with normal karyotype can predict mutations for molecular markers and allows novel insights into perturbed biological pathways. Leukemia 2010;24(6):1216–20 doi 10.1038/leu.2010.73.

47. Bullinger L, Dohner K, Bair E, Frohling S, Schlenk RF, Tibshirani R, et al. Use of gene-expression profiling to identify prognostic subclasses in adult acute myeloid leukemia. N Engl J Med 2004;350(16):1605–16 doi 10.1056/NEJMoa031046.

48. Charlton J, Irtan S, Bergeron C, Pritchard-Jones K. Bilateral Wilms tumour: a review of clinical and molecular features. Expert Rev Mol Med 2017;19:e8 doi 10.1017/erm.2017.8.

49. Taylor MD, Northcott PA, Korshunov A, Remke M, Cho YJ, Clifford SC, et al. Molecular subgroups of medulloblastoma: the current consensus. Acta Neuropathol 2012;123(4):465–72 doi 10.1007/s00401-011-0922-z.

50. Prithviraj P, Anaka M, McKeown SJ, Permezel M, Walkiewicz M, Cebon J, et al. Pregnancy associated plasma protein-A links pregnancy and melanoma progression by promoting cellular migration and invasion. Oncotarget 2015;6(18):15953–65 doi 10.18632/oncotarget.3643.

51. Apps JR, Carreno G, Gonzalez-Meljem JM, Haston S, Guiho R, Cooper JE, et al. Tumour compartment transcriptomics demonstrates the activation of inflammatory and odontogenic programmes in human adamantinomatous craniopharyngioma and identifies the MAPK/ERK pathway as a novel therapeutic target. Acta Neuropathol 2018;135(5):757–77 doi 10.1007/s00401-018-1830-2.

52. Alexandrov LB, Kim J, Haradhvala NJ, Huang MN, Tian Ng AW, Wu Y, et al. The repertoire of mutational signatures in human cancer. Nature 2020;578(7793):94–101 doi 10.1038/s41586-020-1943-3.

53. Alexandrov LB, Nik-Zainal S, Wedge DC, Aparicio SA, Behjati S, Biankin AV, et al. Signatures of mutational processes in human cancer. Nature 2013;500(7463):415–21 doi 10.1038/nature12477.

54. Petljak M, Alexandrov LB, Brammeld JS, Price S, Wedge DC, Grossmann S, et al. Characterizing Mutational Signatures in Human Cancer Cell Lines Reveals Episodic APOBEC Mutagenesis. Cell 2019;176(6):1282–94 e20 doi 10.1016/j.cell.2019.02.012.

55. Papaemmanuil E, Rapado I, Li Y, Potter NE, Wedge DC, Tubio J, et al. RAG-mediated recombination is the predominant driver of oncogenic rearrangement in ETV6-RUNX1 acute lymphoblastic leukemia. Nat Genet 2014;46(2):116–25 doi 10.1038/ng.2874.

56. Zheng S, Cherniack AD, Dewal N, Moffitt RA, Danilova L, Murray BA, et al. Comprehensive Pan-Genomic Characterization of Adrenocortical Carcinoma. Cancer Cell 2016;29(5):723–36 doi 10.1016/j.ccell.2016.04.002.

57. Hoang ML, Chen CH, Sidorenko VS, He J, Dickman KG, Yun BH, et al. Mutational signature of aristolochic acid exposure as revealed by whole-exome sequencing. Sci Transl Med 2013;5(197):197ra02 doi 10.1126/scitranslmed.3006200.

58. Katzenstein HM, Langham MR, Malogolowkin MH, Krailo MD, Towbin AJ, McCarville MB, et al. Minimal adjuvant chemotherapy for children with hepatoblastoma resected at diagnosis (AHEP0731): a Children’s Oncology Group, multicentre, phase 3 trial. Lancet Oncol 2019;20(5):719–27 doi 10.1016/S1470-2045(18)30895-7.

59. Ruggiero A, Trombatore G, Triarico S, Arena R, Ferrara P, Scalzone M, et al. Platinum compounds in children with cancer: toxicity and clinical management. Anticancer Drugs 2013;24(10):1007–19 doi 10.1097/CAD.0b013e3283650bda.

60. Smith KS, Xu K, Mercer KS, Boop F, Klimo P, DeCupyere M, et al. Patient-derived orthotopic xenografts of pediatric brain tumors: a St. Jude resource. Acta Neuropathol 2020;140(2):209–25 doi 10.1007/s00401-020-02171-5.

61. CCDI Symposium Childhood Cancer. <https://www.cancer.gov/news-events/cancer-currents-blog/2019/lowy-ccdi-symposium-childhood-cancer>.

62. Li H, Handsaker B, Wysoker A, Fennell T, Ruan J, Homer N, et al. The Sequence Alignment/Map format and SAMtools. Bioinformatics 2009;25(16):2078–9 doi 10.1093/bioinformatics/btp352.

63. Hankins JS, Estepp JH, Hodges JR, Villavicencio MA, Robison LL, Weiss MJ, et al. Sickle Cell Clinical Research and Intervention Program (SCCRIP): A lifespan cohort study for sickle cell disease progression from the pediatric stage into adulthood. Pediatr Blood Cancer 2018;65(9):e27228 doi 10.1002/pbc.27228.

64. Li H. Aligning sequence reads, clone sequences and assembly contigs with BWA-MEM. arXiv: 2013;1303.3997v1

65. Ryan Poplin VR-R, Mark A. DePristo, Tim J. Fennell, Mauricio O. Carneiro, Geraldine A. Van der Auwera, David E. Kling, Laura D. Gauthier, Ami Levy-Moonshine, David Roazen, Khalid Shakir, Joel Thibault, Sheila Chandran, Chris Whelan, Monkol Lek, Stacey Gabriel, Mark J. Daly, Benjamin Neale, Daniel G. MacArthur, and Eric Banks. Scaling accurate genetic variant discovery to tens of thousands of samples. bioRxv 2017.

66. Microsoft. <https://azure.microsoft.com/mediahandler/files/resourcefiles/accelerate-precision-medicine-with-microsoft-genomics/Accelerate_precision_medicine_with_Microsoft_Genomics.pdf>.

67. Picard. <http://broadinstitute.github.io/picard/>.

68. FASTQC. <https://www.bioinformatics.babraham.ac.uk/projects/fastqc/>.

69. Okonechnikov K, Conesa A, Garcia-Alcalde F. Qualimap 2: advanced multi-sample quality control for high-throughput sequencing data. Bioinformatics 2016;32(2):292–4 doi 10.1093/bioinformatics/btv566.

70. QC workflow. < https://github.com/stjudecloud/rfcs/blob/rfcs/qc-workflow/text/0002-quality-check-workflow.md>.

71. Rnaseq workflow v2.0.0. <https://stjudecloud.github.io/rfcs/0001-rnaseq-workflow-v2.0.0.html>.

72. Dobin A, Davis CA, Schlesinger F, Drenkow J, Zaleski C, Jha S, et al. STAR: ultrafast universal RNA-seq aligner. Bioinformatics 2013;29(1):15–21 doi 10.1093/bioinformatics/bts635.

73. Gencode v31. <https://www.gencodegenes.org/human/release_31.html >.

74. Anders S, Pyl PT, Huber W. HTSeq--a Python framework to work with high-throughput sequencing data. Bioinformatics 2015;31(2):166–9 doi 10.1093/bioinformatics/btu638.

75. Ngsderive. <https://github.com/stjudecloud/ngsderive>.

76. Zhang J, Ding L, Holmfeldt L, Wu G, Heatley SL, Payne-Turner D, et al. The genetic basis of early T-cell precursor acute lymphoblastic leukaemia. Nature 2012;481(7380):157–63 doi 10.1038/nature10725.

77. Chen X, Gupta P, Wang J, Nakitandwe J, Roberts K, Dalton JD, et al. CONSERTING: integrating copy-number analysis with structural-variation detection. Nat Methods 2015;12(6):527–30 doi 10.1038/nmeth.3394.

78. Alexandrov LB, Nik-Zainal S, Wedge DC, Campbell PJ, Stratton MR. Deciphering signatures of mutational processes operative in human cancer. Cell Rep 2013;3(1):246–59 doi 10.1016/j.celrep.2012.12.008.

