## Supplementary Data for "St. Jude Cloud—a Pediatric Cancer Genomic Data Sharing Ecosystem"

* Contributed Equally

### SUPPLEMENTARY TABLES

Supplementary Table S1. Data Sets in PeCan Knowledgebase.

Supplementary Table S2. **(A)** Gene set enrichment analysis following differential gene expression analysis between two osteosarcoma groups (OS group1 and OS group 2) identified in **Fig. 5C**. Table displays Human Gene Atlas gene groups over-represented in upregulated genes in OS group 2 following ENRICHR analysis. **(B)** Gene set enrichment analysis following differential gene expression analysis between two adamantinomatous craniopharyngioma groups (ACPG group1 and ACPG group 2) identified in **Fig. 5E**. Table displays Human Gene Atlas gene groups over-represented in upregulated genes in ACPG group 1 following ENRICHR analysis. **(C)** Gene set enrichment analysis following differential gene expression analysis between two adamantinomatous craniopharyngioma groups (ACPG group1 and ACPG group 2) identified in **Fig. 5E**. Table displays WikiPathways 2019 Human gene groups over-represented in upregulated genes in ACPG group 1 following ENRICHR analysis. **(D)** Gene set enrichment analysis following differential gene expression analysis between two adamantinomatous craniopharyngioma groups (ACPG group1 and ACPG group 2) identified in **Fig. 5E**. Table displays GO Molecular Function 2018 gene groups over-represented in upregulated genes in ACPG group 2 following ENRICHR analysis.

Supplementary Table S3. Displaying the grouping of different pediatric cancer subtypes into blood, solid, and brain tumor categories in addition to their abbreviations.

### SUPPLEMENTARY FIGURES

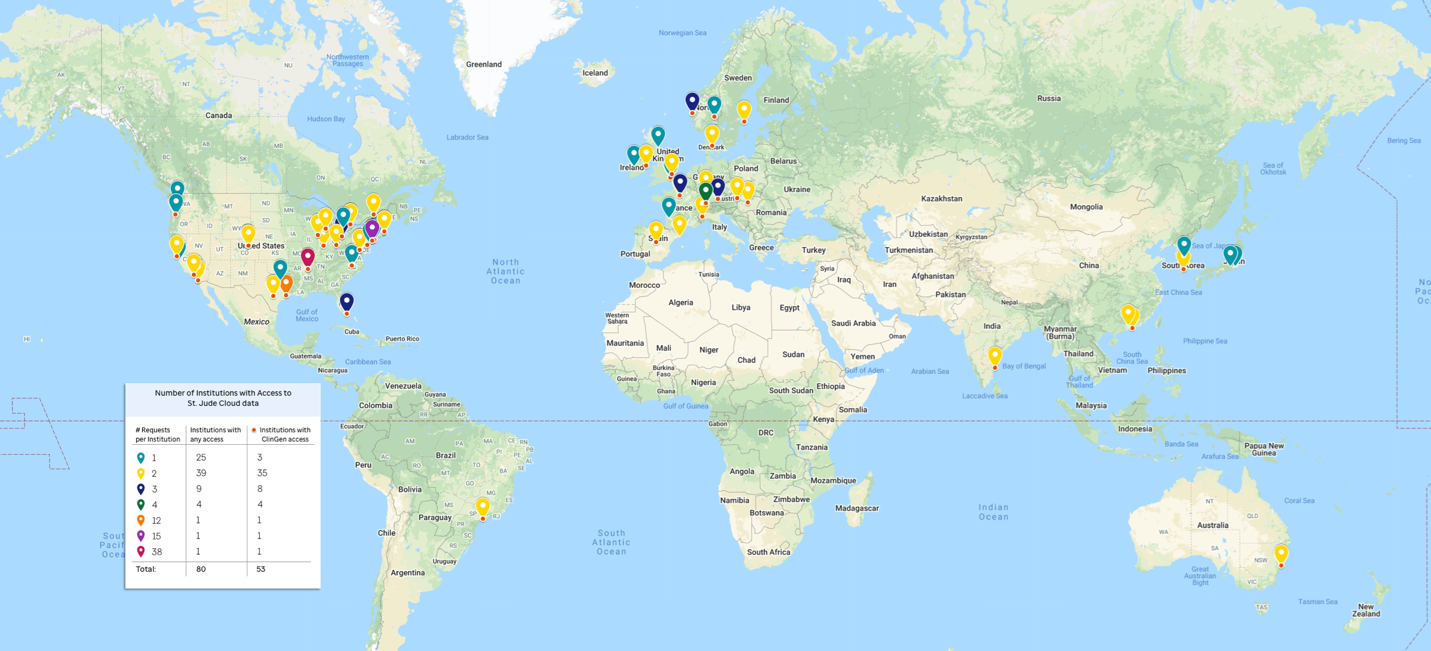

Supplementary Figure S1**. Data sharing with the global research community.** The number of approved data requests per institution throughout the world are indicated by color. The institutions granted access to clinical genomics (ClinGen) data are indicated with a red dot.

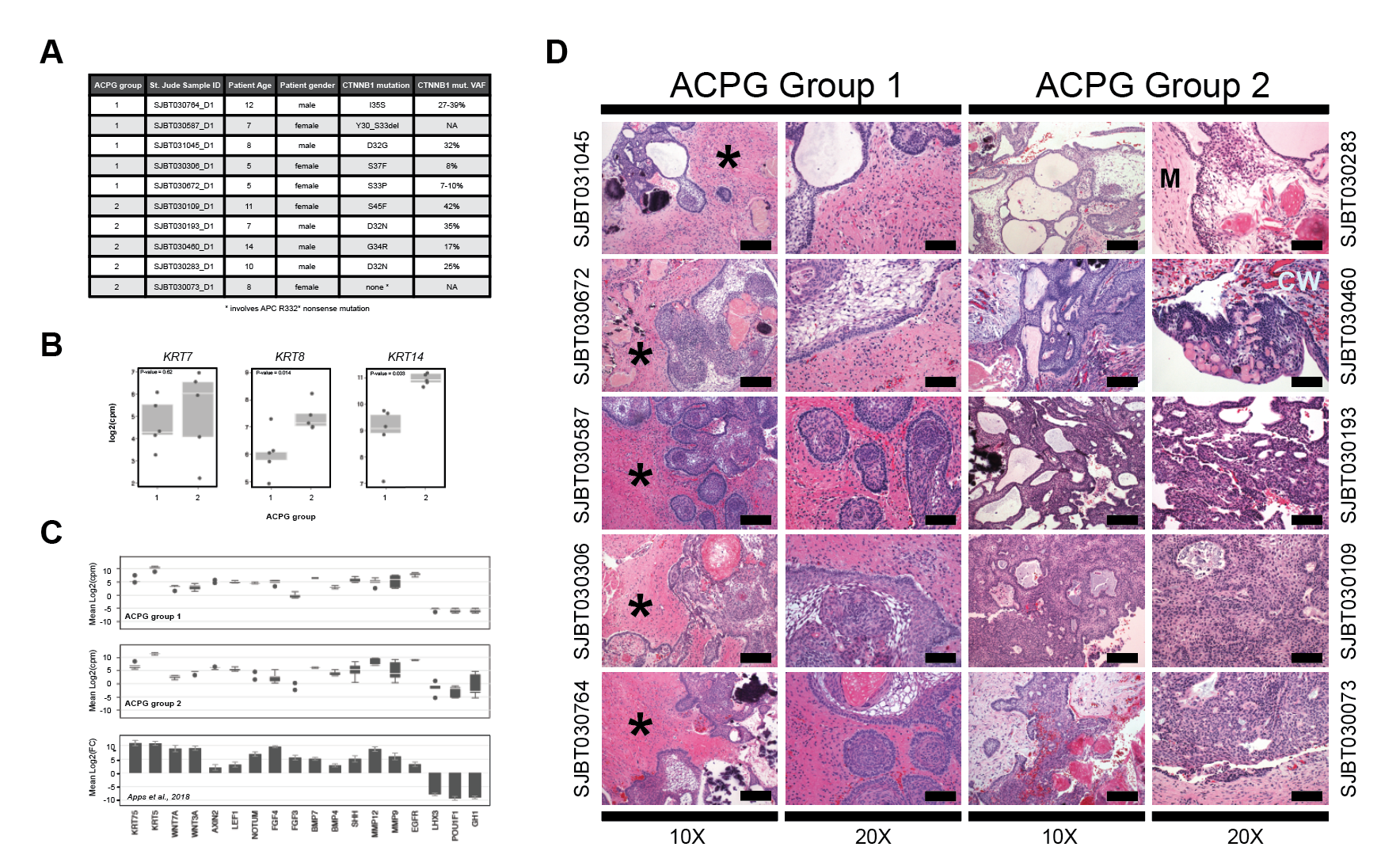

Supplementary Figure S2. **(**A) St. Jude Cloud Adamantinomatous Craniopharyngioma (ACPG) select demographic and genetic attributes. The ACPG Group as defined in Figure 5 is indicated. CTNNB1 mutational variant allele frequencies (VAF) were determined from whole-genome (WGS) or whole-exome (WES) sequencing data. Where CTNNB1 VAFs were calculated from both WGS and WES, a range is presented. (B) Boxplots of Keratin (KRT7, KRT8, and KRT14) gene expression observed in ACPG Group 1 and 2 samples. (C) Boxplots of select absolute gene expression observed in ACPG Group 1 (top) and 2 (middle), in addition to reported differential gene expression for ACPG samples when compared to normal fetal brain tissue (1) (bottom). Note p-values calculated using a t-test. (D) Hematoxylin and Eosin stained sections from ACPG Group 1 and Group 2 samples at 10X and 20X magnification. All five Group 1 tumors were characterized by brain invasion. The surrounding reactive brain parenchyma is designated by an asterisk (*). In contrast, Group 2 tumors showed involvement with either tumor cyst walls (“CW”) or meninges (“M”), with minimal or no involvement with neural tissue. Scale bar represents 160 uM for the 10X images and 80 uM for the 20X images.

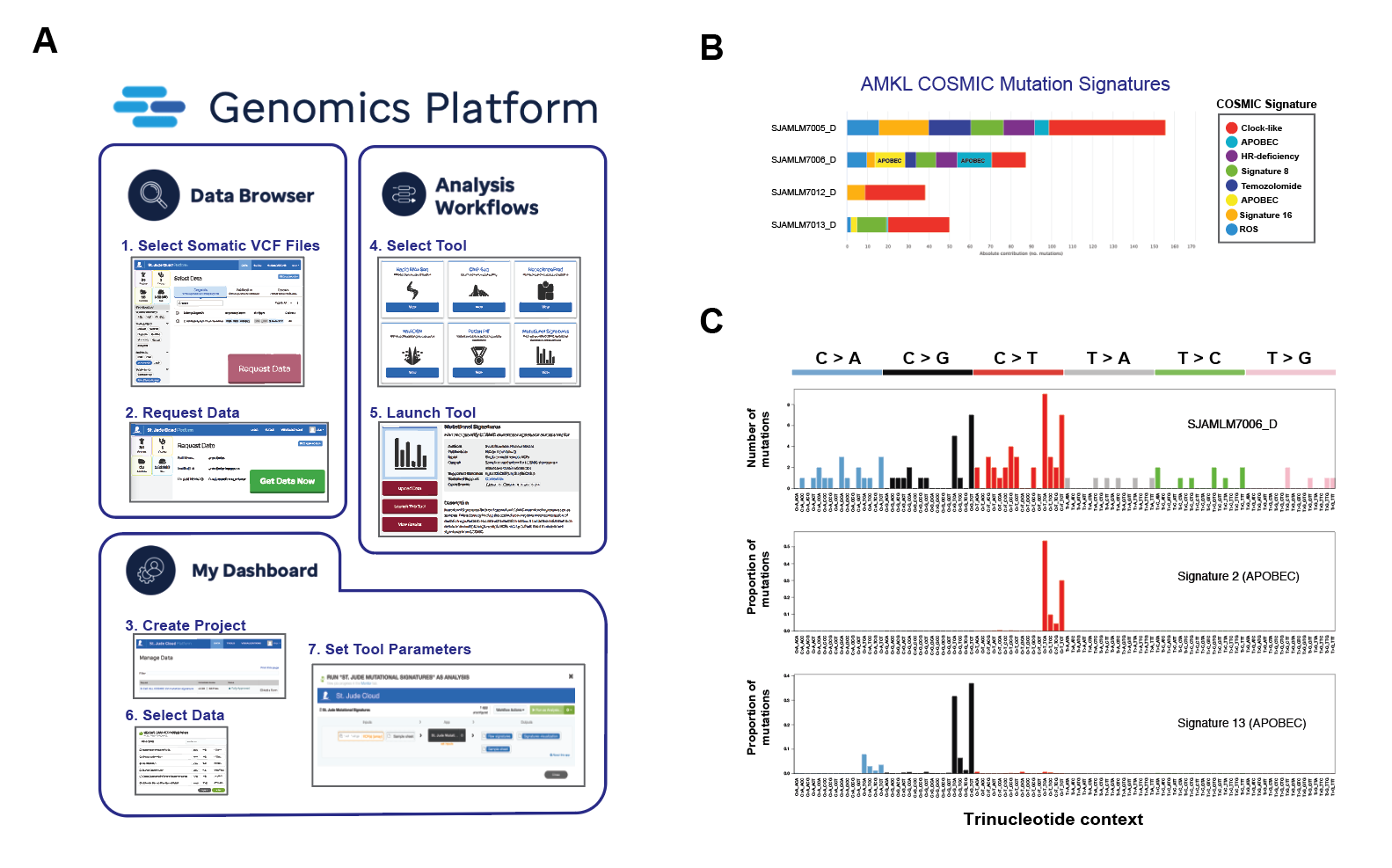

Supplementary Figure S3. Identification of COSMIC mutation signatures in pediatric tumor samples. (A) Step-by-step depiction of the use of St. Jude Cloud Mutational Signatures tool via the St. Jude Cloud *Genomics Platform*. Comprising the i) selection of somatic VCF files, and ii) submission of data access request via the *Data Browser*; iii) creation of a project within a private user workspace accessible via *My Dashboard*; iv) Mutational Signatures tool selection and launch via the *Analysis Workflows*; and iv-vii) running of the tool. (B) Identification of COSMIC mutation signatures within four acute megakaryoblastic leukemias (AMKL) via St. Jude Cloud Mutational Signatures analysis of associated whole-genome sequencing somatic mutations. APOBEC COSMIC mutation signatures (signature 2 and 13) identified in SJAMLM7006_D are indicated. COSMIC mutation signature color key (right). (C) Whole-genome sequencing somatic mutation signature landscape of acute megakaryoblastic leukemia sample SJAMLM7006_D harboring APOBEC COSMIC mutational signatures 2 and 13.

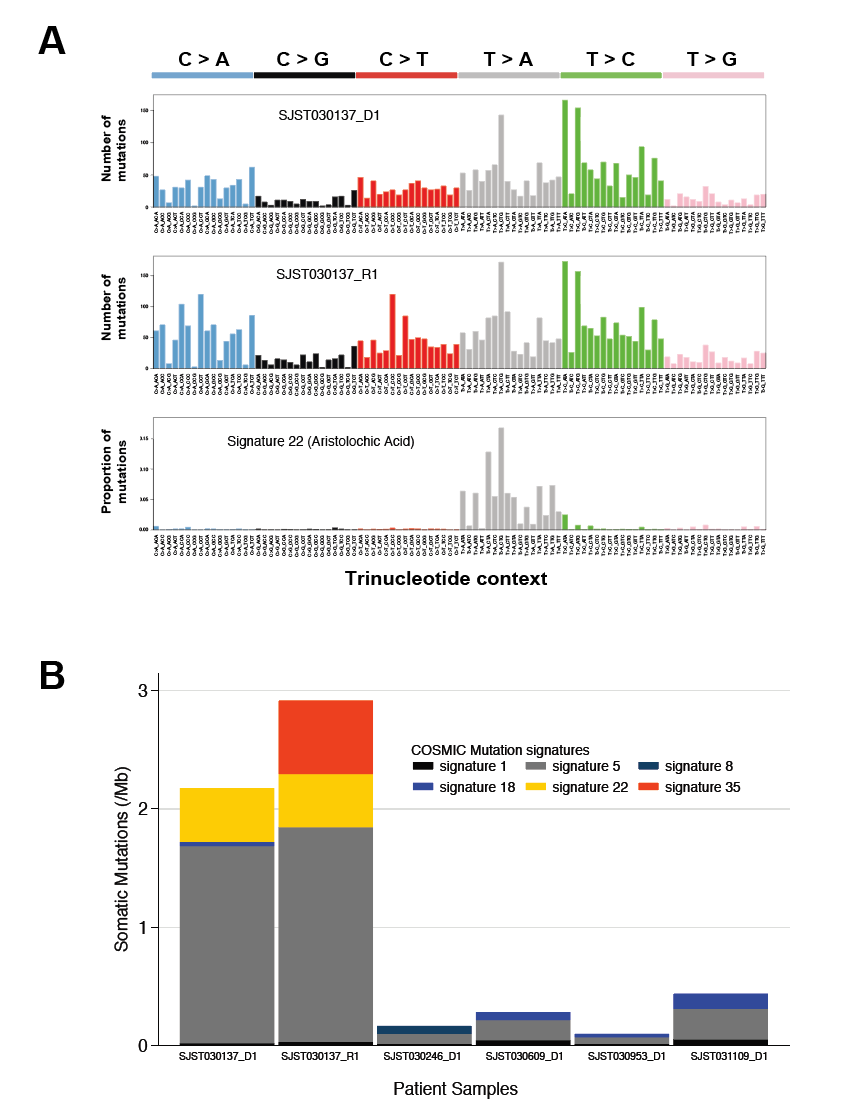

Supplementary Figure S4. (A) Whole-genome sequencing somatic mutation signature landscape of hepatoblastoma patient diagnosis and relapse samples (SJST030137_D1 and SJST030137_R1) harboring Aristolochic Acid COSMIC mutational signature 22. (B) Stacked barplot of somatic mutations (No./Mb) contributing to COSMIC Mutation Signature 1, 5, 8, 18, 22, and 35 within six hepatoblastoma patient samples. Patient SJST030137 involves both diagnostic and relapse samples (SJST030137_D1 and SJST030137_R1), while remaining patients only involve diagnostic samples. Patient SJST030137 is an Asian male while the remaining samples are from non-Asian patients. Data from published (SJST030137_D1, SJST030246_D1, (2)) and unpublished real-time clinical genomics (SJST030137_R1, SJST030609_D1, SJST030953_D1, SJST031109_D1) datasets.

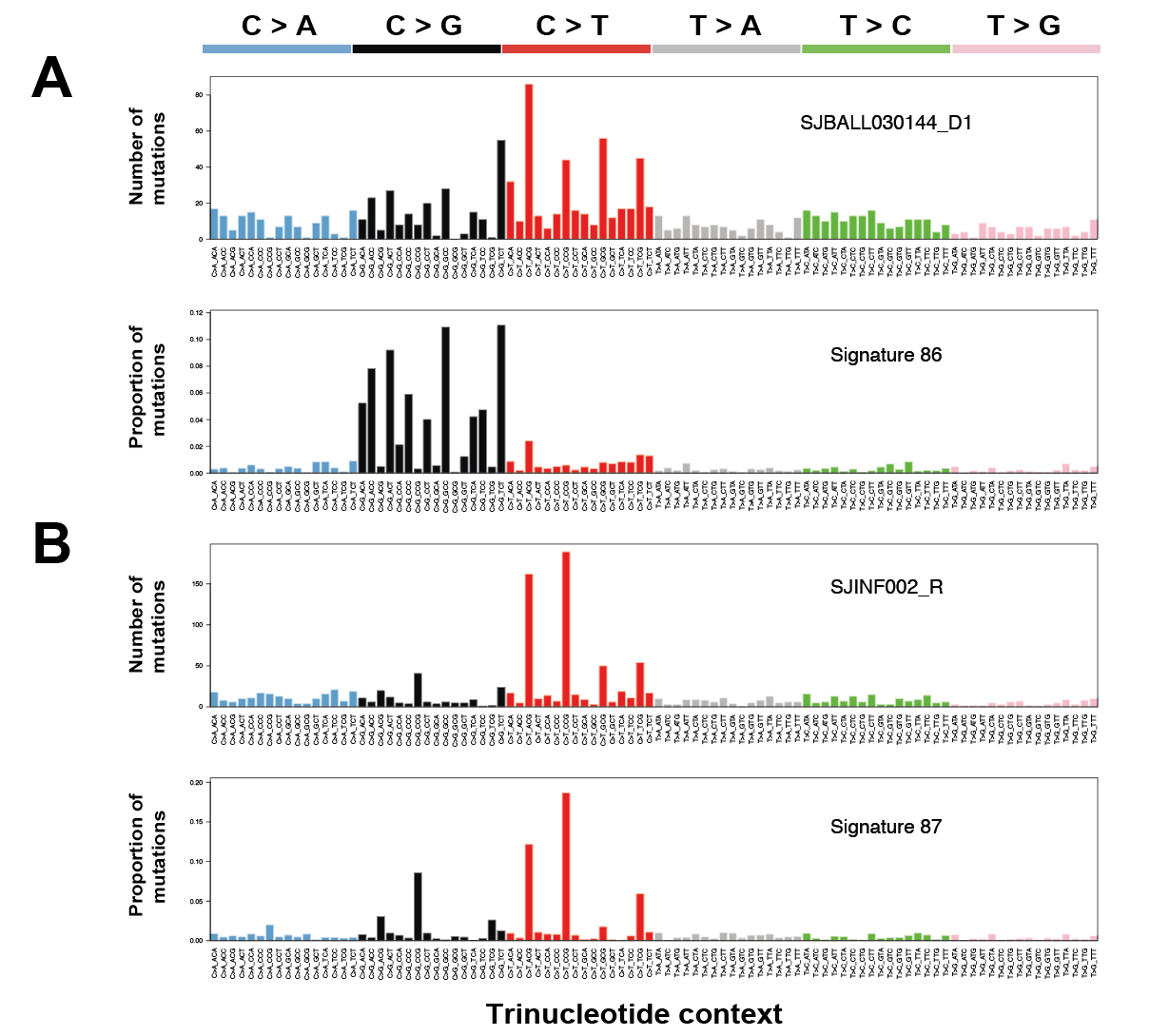

Supplementary Figure S5. (A) Whole-genome sequencing somatic mutation signature landscape of relapsed B-cell acute lymphoblastic leukemia sample harboring therapy-related COSMIC mutational signature 86. (B) Whole-genome sequencing somatic mutation signature landscape of relapsed B-cell acute lymphoblastic leukemia sample harboring therapy-related COSMIC mutational signature 87.

### REFERENCES

1.	Apps JR, Carreno G, Gonzalez-Meljem JM, Haston S, Guiho R, Cooper JE*, et al.* Tumour compartment transcriptomics demonstrates the activation of inflammatory and odontogenic programmes in human adamantinomatous craniopharyngioma and identifies the MAPK/ERK pathway as a novel therapeutic target. Acta Neuropathol **2018**;135(5):757-77 doi 10.1007/s00401-018-1830-2.

2.	Genome4Kids.  <<https://clinicaltrials.gov/ct2/show/NCT02530658>>.
